# ImmunoCluster: A computational framework for the non-specialist to profile cellular heterogeneity in cytometry datasets

**DOI:** 10.1101/2020.09.09.289033

**Authors:** James W. Opzoomer, Jessica Timms, Kevin Blighe, Thanos P. Mourikis, Nicolas Chapuis, Richard Bekoe, Sedigeh Kareemaghay, Paola Nocerino, Benedetta Apollonio, Alan G. Ramsay, Mahvash Tavassoli, Claire Harrison, Francesca Ciccarelli, Peter Parker, Michaela Fontenay, Paul R. Barber, James N. Arnold, Shahram Kordasti

## Abstract

High dimensional cytometry is an innovative tool for immune monitoring in health and disease, it has provided novel insight into the underlying biology as well as biomarkers for a variety of diseases. However, the analysis of multiparametric “big data” usually requires specialist computational knowledge. Here we describe *ImmunoCluster (https://github.com/kordastilab/ImmunoCluster)* an R package for immune profiling cellular heterogeneity in high dimensional liquid and imaging mass cytometry, and flow cytometry data, designed to facilitate computational analysis by a non-specialist. The analysis framework implemented within *ImmunoCluster* is readily scalable to millions of cells and provides a variety of visualization and analytical approaches, as well as a rich array of plotting tools that can be tailored to users’ needs. The protocol consists of three core computational stages: 1, data import and quality control, 2, dimensionality reduction and unsupervised clustering; and 3, annotation and differential testing, all contained within an R-based open-source framework.

## Introduction

Systems immunology approaches aim to explore and understand the complexity of the immune system. However, with 350 CD (‘cluster of differentiation’) antigens, over 100 cytokines and chemokines, and many different cell subsets this is a challenging task [1]. Liquid mass cytometry (LMC), imaging mass cytometry (IMC) and flow cytometry (FC) are powerful techniques applied to exploratory immunophenotyping and biomarker discovery, with the ability to profile over 40 markers on an individual cell these techniques have rapidly expanded our understanding of the immune system and its perturbations throughout disease pathogenesis [2, 3]. However, such comprehensive analyses produce large amounts of data and increased dimensionality, resulting in a demanding computational task to analyze [4]. Often, a limiting factor to these analyses are the requirement of an in-depth knowledge of computational biology. There is an unmet need for an easy-to-use open-source computational framework, to explore high dimensional single-cell cytometry datasets such as LMC, IMC and FC for non-specialists. Thus, we developed a framework which will provide appropriate data structures and methods to increase the utility of these techniques and make data analysis and interpretation accessible to all researchers to facilitate immune phenotyping projects, such as those associated with clinical trials.

Here, we present *ImmunoCluster*, an open-source computational framework for the analysis of high-dimensional LMC, IMC, and FC datasets. *ImmunoCluster* is an R package and framework designed for discovery, visualization of immune subpopulations, and analysis of changes in cell phenotypes and abundance across experimental conditions, including serial samples from a clinical trial setting. The computational approaches used to analyze high dimensional data are rapidly evolving and *ImmunoCluster* has the flexibility to incorporate these novel computational methods in dimensionality reduction, clustering and trajectory analysis within its framework. The computational framework is designed for ease-of-use and broad applicability, so that it is simple enough to be implemented by users with only a basic knowledge of R, as well as possessing the flexibility for more advanced users to build on the framework. Our framework relies on the SingleCellExperiment class [5], this means that the Flow Cytometry Standard (FCS) data is contained within a purpose-built object that stores all stages of analysis to permit multiple analysis paths to be performed in parallel, along with convenient ‘wrapper functions’ allowing for the use of popular dimensionality reduction and clustering algorithms in an easy format for non-specialists to rapidly produce interpretable data that are extendable to advanced experimental designs. The outputs from *ImmunoCluster* are created using the *ggplot* package which generates high-resolution images that can be utilized directly into figures for reports and publications. Importantly, and unique to *ImmunoCluster*, the scope of functionality in this package also enables the analysis of IMC data with minimal supervision. IMC provides a means to analyze the spatial dimension of the cells *in situ* within tissues, which can provide important insight when ascribing functionality to cell populations. Methods for analyzing large cytometry datasets, in particular IMC datasets, in an open-source computational environment are currently limited. *ImmunoCluster* has been designed for use by a non-specialist, however would be useful to a range of users from wet-lab non-specialist experimentalists to experienced computational biologists with potential utility in day-to-day research through to large-scale clinical trials involving multiple longitudinal serial biopsies where samples need to be analyzed and compared.

Clinical immune monitoring is an area of increasing utilization [6]. In the current work we will present examples of *ImmunoCluster’s* ability to detect clinically relevant perturbations in immune cellular heterogeneity in LMC, IMC and FC datasets in human patient and/or healthy samples. *ImmunoCluster* is easily extensible to several arbitrary experimental designs and can be utilized to aid both discovery of novel subpopulations of cells within heterogeneous samples or as a foundation for monitoring longitudinal changes or responses to treatment. As such, *ImmunoCluster* provides a resource that will permit high dimensional data to be more widely adopted in immuno-biology and many other disciplines.

## Methods

### Data file pre-processing and preparation for implementation into the *ImmunoCluster* framework Liquid mass cytometry data

Here we describe an example of the *ImmunoCluster* package analyzing previously published mass cytometry data (CyTOF^®^) from 15 leukemia patients 30 and 90 days after bone marrow transplantation (BMT) which has been published by Hartmann *et al.* [7]. After BMT three of these patients suffered acute graft versus host disease (GvHD) and 12 had no evidence of GvHD (noted as ‘none’ in all figures). For systems-level biomarker discovery within this trial, a panel of 33 antibodies were incorporated into the immunophenotyping panel (Table S1). The panel was designed to cover all major immune cell lineages and several functional subsets including; T-, B-, NK and myeloid cells and granulocytes (detailed in [7]). The marker panel also included a variety of immune-regulatory proteins, such as PD-1, PD-L1, TIM3, and TCRVa24-Ja18. The publicly available dataset from the Hartmann *et al.* study was extracted for this study from the FlowRepository (http://flowrepository.org/id/FR-FCM-Z244) [8]. Raw FCS files for the LMC dataset collected on the Helios™ CyTOF^®^ system (Fluidigm, UK) machine were first normalized using the free CyTOF^®^ 6.7 system control software (files can also be concatenated using the same software if necessary). These files were gated using FlowJo™ (version: 10.5.3) (Becton, Dickinson and Company, UK) to remove beads and doublets, dead and non-CD45^+^ cells as well as erythrocytes (CD235αβ/CD61^+^) and neutrophils (CD16^+^) (Figure S1), as described by Hartmann *et al.* [7], before implementation into the *ImmunoCluster* framework. There are various alternative open-access tools which can be used to gate these files, such as OpenCyto [9], CytoExploreR [10] and Cytoverse (https://cytoverse.org/).

### Imaging mass cytometry data

Two different examples of IMC datasets were implemented into the framework. One IMC dataset was from a human tissue section taken from a patient with head and neck squamous cell carcinoma (HNSCC). Consent was attained by the King Guy’s & St Thomas’ Research Biobank, within King’s Health Partners Integrated Cancer Centre. The second IMC dataset was from a lymph node section from a patient with diffuse large B-cell lymphoma (DLBCL). Formalin-fixed paraffin-embedded (FFPE) DLBCL tumor tissue was obtained from King’s College Hospital, in accordance with the Declaration of Helsinki and approved by the UK National Research Ethics Committee (reference 13/NW/0040). A detailed protocol is available for both datasets as a Supplementary File (File S1 and S2, respectively). The raw HNSCC IMC data from 12 channels (10 markers + 2 intercalator-Ir channels (Table S2), and the raw DLBCL IMC data from 20 channels (19 markers + nuclei channel) (Table S3), was processed with Python scripts and *CellProfiler* pipelines according to Schulz *et al.* [11] to segment the image data into individual cells. The mean marker intensity for each segmented cell was exported (comma-separated values (CSV) file) and divided into 6 regions of equal area (3 tumor and 3 stroma regions) (Figure S2). The mean marker intensity for each segmented cell was multiplied by 65535 to recover the image intensities (Fluidigm IMC machine = 16-bit, the dynamic range of the measurements are 0 to (2^16^)-1) and asinh (co-factor 0.8) transformed in R Studio. These CSV files were converted into FCS files and gated (Figure S2) in Cytobank [12], a function is also available within the *ImmunoCluster* package to convert files into FCS format.

### Flow cytometry data

FC data from the BM of 7 healthy donors (HDs) were used to demonstrate the ability of the framework to analyze conventional FC data and identify rare immune cell populations. The BM samples from elderly HDs were obtained by extraction of the bone marrow cells from the bone of the femoral head. This non-interventional study was approved by the ethical committee of Cochin-Port Royal Hospital (Paris, France) (CLEP Decision N°: REC number). Femoral heads were obtained after informed consent, during hip replacement surgery. These were cut in half and collected in a conservation medium (Hanks balanced salt solution with NaHCO_3_, Eurobio ™), supplemented with heparin (7%) and then transported to the laboratory at room temperature (RT). These were scraped with a spatula, ground in a mortar, and washed with a PBS solution supplemented with DNAse (Sigma Aldrich) at 100 ug/ml. The FCS files were gated for singlets using a FSCint/FSCpeak dot plot, and dead cells were eliminated using a FSC/SSC dot plot. Leukocytes were gated on a CD45/SSC dot plot. Finally, they were gated to identify the CD3^+^ CD4^+^ population of cells for analysis in the *ImmunoCluster* framework (gated in Cytobank [12]).

### Accessing the open-source *ImmunoCluster* tool

The *ImmunoCluster* package is accessible via GitHub (https://github.com/kordastilab/ImmunoCluster), where a detailed step-by-step walkthrough of the tool is available. A number of previous workflow pipelines influenced the development of *ImmunoCluster*, including *CyTOF workflow* [13], *CATALYST* [14], *cytoClustR* [15], and Diggins *et al.* methodology [16]. The framework has incorporated methods for applying popular R packages (i.e. *RPhenograph*, *Rtsne* and *FlowSOM*), and its visualization tools were developed using *scDataviz*, a Bioconductor package [17] for visualizing single cell data. Users will need to have R downloaded onto their desktop to run the *ImmunoCluster* tool.

### Workflow overview

The *ImmunoCluster* framework provides tools and support to allow researchers to follow a workflow which guides them through experimental design, data analyses and interpretation, to publishable graphics identifying differences in phenotype and abundance of cells between conditions analyzed (Figure 1 and 2). The framework comprises three core computational stages which are conducted by the *ImmunoCluster* tool:

**Stage 1. Data import and quality control**

a. We provide tools designed primarily for computational pre-processing of FCS data file for downstream applications including parameter renaming and transformation. The generated high dimensional data are imported into the computational framework as FCS file formats or summarized feature expression values in CSV tabular format.
b. Associated metadata files need to be completed by the researcher in the experimental design stage (Figure 1). This includes a sample metadata file, in which all metadata relating to samples will need to be input, such as timepoints, response to treatments and patient status (Figure S3A). For simplicity in executing this step, the panel metadata file allows users to rename parameters, such as the markers used in the analysis as well as select the markers which are to be used for different analyses stages (all markers may not need to be included in the following steps) (Figure S3B).
c. All of the above mentioned data is stored within a SingleCellExperiment object [5]. The SingleCellExperiment is an S4 class object from Bioconductor that is implemented into the *ImmunoCluster* framework, and is in essence a data container in which you can store and retrieve information such as metal-barcoding for sample multiplexing, metadata, dimensionality reduction coordinates and more (outlined in Figure 1).
d. After the data and metadata files have been imported and stored in the SingleCellExperiment object, we provide several approaches for initial exploratory visualization of the data. One example of this is the Multidimensional scaling (MDS) plots, these allow for an initial visualization of data points (labeled with metadata) and show the similarity and differences of the samples in two-dimensions which can be used to detect batch effects and other technical artifacts such as antibody staining anomalies. Additionally, a heatmap of expression of markers measured for each patient can be created and metadata can be used to annotate the heatmap.
**Stage 2. Dimensionality reduction and unsupervised clustering**
 
a. Dimensionality reduction is a key component of high-dimensional single-cell data analyses and enables researchers to view high-dimensional data. For example, the 33 protein markers analyzed by Hartmann *et al.* [7] can be reduced to lower dimensional embedded coordinates per cell. Nonlinear dimensionality reduction techniques can avoid overcrowding and represent data in distinct cell islands (Figure 2).
b. The framework offers users three nonlinear statistical methods for representing high dimensional single-cell data in low-dimensional space (Figure S4). MDS (utilized in stage 1 for initial data exploration), uniform manifold approximation and projection (*UMAP*) [18], and t-Distributed Stochastic Neighbor Embedding (*tSNE*) [19] algorithms are available to generate a 2 dimension embedding of the data. However, the SingleCellExperiment object can store any other form of arbitrary dimensionality reduction, such as principle component analysis (PCA) [20] or DiffusionMaps [21], amongst others.
c. After the data has been visualized in dimensionality reduced space, clustering algorithms can be used to identify cell communities within the data, which allows the identification of phenotype and abundance of cell clusters within populations/groups being analyzed. A number of clustering algorithms are available within the framework to define cell populations of interest in an unbiased manner. Firstly, an ensemble (use of multiple algorithms) clustering method of *FlowSOM* [22] and Consensus clustering (*ConsensusClusterPlus* R package [23]), which tests the stability of the clusters, leading to better results than applying a basic hierarchical clustering algorithm. Additionally, the *PhenoGraph* clustering algorithm is available, this uses *K*-nearest neighbors and a Euclidean distance metric [24]. Either or both clustering algorithms can be applied at the users’ discretion.
d. The aim of these clustering algorithms is to assign all cells to *K* clusters (*K*_1_, *K*_*2*_,…) resulting in clusters corresponding to true cell types. Typically, a *K* larger than the number of expected cell types is chosen at this stage as the *ImmunoCluster* framework allows users to explore all *K* clusters (Figure S5), as well as collapsing clusters of the same cell type into one cluster if the user feels over clustering has occurred. Over clustering may enable the clustering identification of rare cell types, sometimes at the expense of creating several ‘artificial’ clusters of the more prevalent cell types. Additionally, the workflow allows the generation of an elbow plot which can also be created to help the user to select an appropriate number of clusters (Figure S6).
**Stage 3. Annotation and differential testing**

a. After dimensionality reduction and clustering the next step is to annotate each cluster based on marker expression. If over-clustering has occurred, multiple clusters of the same cell type can be present, and either a lower number of *K* clusters can be used, additionally the clusters of the same cell type can be collapsed into each other to create one cluster.
b. Clusters containing multiple cell types may also be identified, this could mean the user has under-clustered, or occasionally a group of different cells are clustered together by the algorithms.
c. Expression of each marker analyzed can be projected onto the dimensionality reduced data by the researcher, aiding with the identification of cell types or phenotypically distinct clusters.
d. Outputs such as heatmaps showing the expression of all markers for each cluster can be used to identify the clusters of interest (Figure S7). These can be used to annotate figures produced in Stage 3.
e. Due to the type of data produced from IMC (lower resolution compared to LMC and FC) a rank heatmap is created in which the expression of each marker is ranked. This rank heatmap can be used to identify clusters which are ‘high’ or ‘low’ for particular markers to distinguish and logically assign cell populations manually.
f. These tools help define biological identity of cell clusters from Stage 2 and construct a hierarchy of biologically meaningful cell populations based on marker expression and similarity.
g. The metadata added to the SingleCellExperiment can be used to annotate the dimensionality reduced (i.e. *UMAP*) and clustered (*FlowSOM* or *Phenograph*) data, allowing for visualization of the distribution of cell islands between different condition/timepoints (Figure 2).
h. The above-mentioned metadata is also used to annotate all differential testing outputs, meaning the user can make many comparisons and explore multiple parameters all within the *ImmunoCluster* framework.
i. A number of differential testing outputs can be created, such as median marker expression for each sample (using clustered data) can be used to produce a hierarchical clustered heatmap, box plots of cell cluster abundance between different conditions at different time points (any metadata can be used to annotate these figures).
j. Statistically significant differences in the annotated populations between conditions are identified and displayed throughout the framework to the user for ongoing investigation and interpretation. In IMC, positional data can be used to overlay cell location of populations to investigate co-localization and density maps.

**Figure 1.**
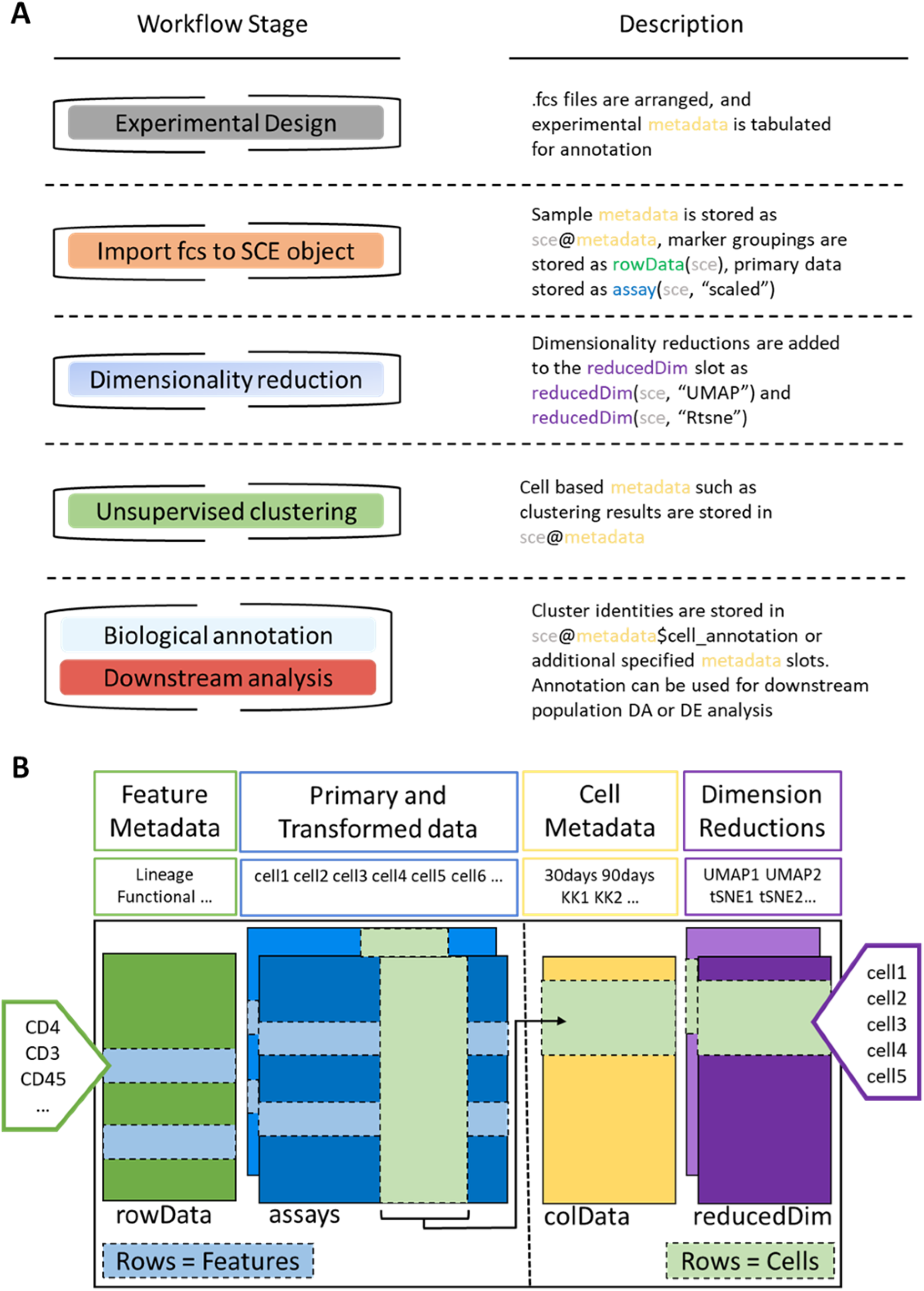
*ImmunoCluster* workflow stages and SingleCellExperiment structure. **A)** Schematic outlining the typical mass cytometry workflow and its interactions with the SingleCellExperiement *ImmunoCluster* object. **B)** SingleCellExperiment structure in *ImmunoCluster*. The SingleCellExperiment class is a data container, storing multiple layers of data to create the SingleCellExperiment object which holds all relevant data for an experiment. **Feature Metadata**: imported by the user in the form of a panel metadata file, which is a table containing all markers measured, each annotated with either lineage or functional information for downstream analysis. **Primary and Transformed data:** the imported expression data is stored in an assay, additionally the scaled data (arcsinh transformed) is also stored in an assay, meaning both can be easily accessed. **Cell Metadata:** the first metadata added to this element of the structure will be a sample metadata file imported by the user, containing any relevant metadata for the experiment i.e., days after treatment and GvHD or non-GvHD. Throughout the *ImmunoCluster* tool more layers of metadata are added to Cell Metadata, i.e., cell cluster identification (*FlowSOM* and *Phenograph*). **Dimension Reductions:** dimensionality reduction coordinates, such as *UMAP* and *tSNE* are stored and can be easily accessed throughout the *ImmunoCluster* tool for downstream analyses.

**Figure 2.**
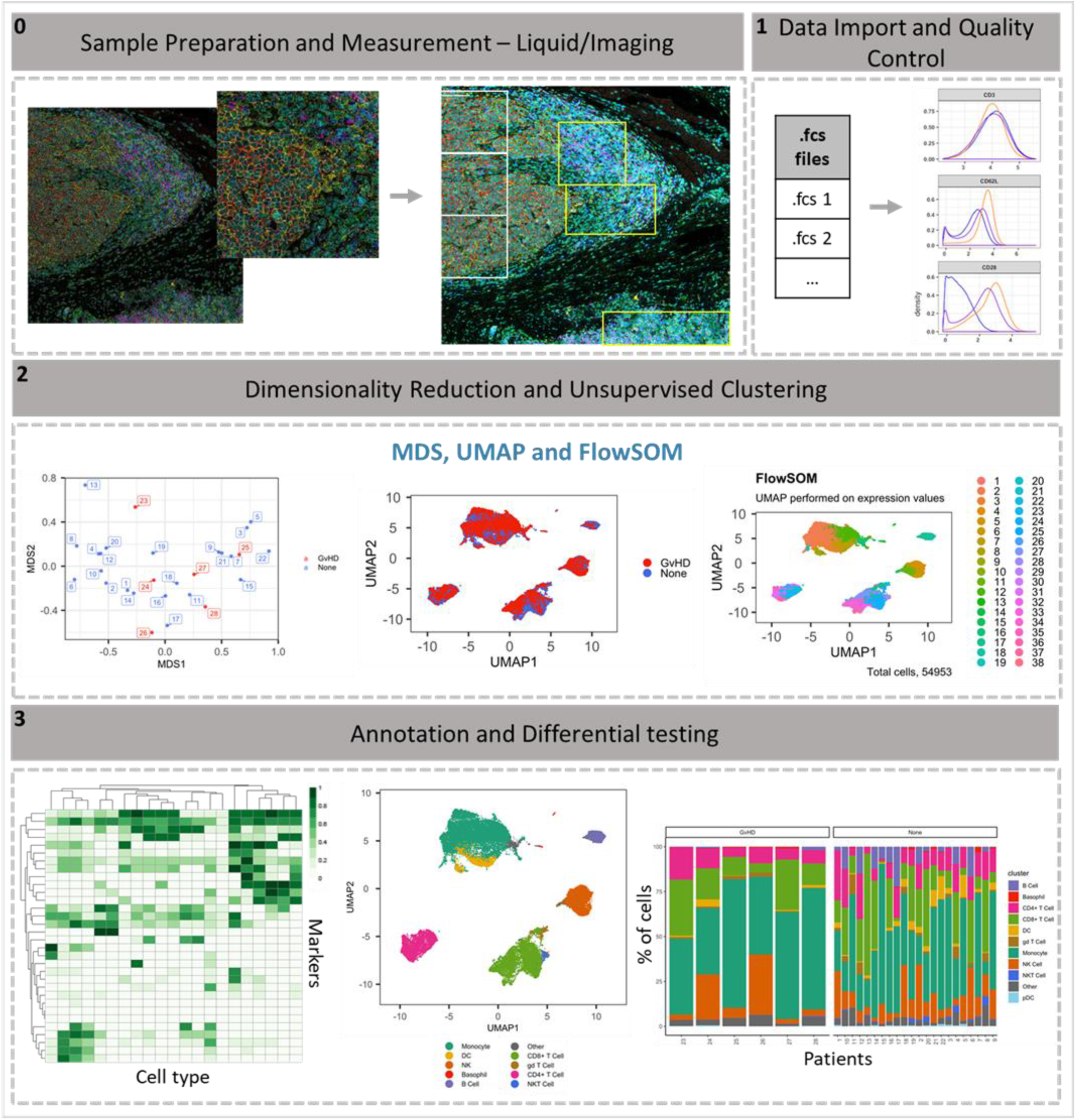
*ImmunoCluster* workflow overview. **0)** Samples are stained/treated and measured, for IMC the tissue is segmented, and regions selected for further downstream analysis. After measurement the raw data is normalized, concatenated (combining FCS files from the same samples which may have been split due to large sample volume or technical issues) and gated, before importing into *ImmunoCluster*. **1)** Quality control of data is carried out before analysis. **2)** Data is reduced to two dimensions using either *UMAP* or *tSNE* algorithm, and data are clustered using the *FlowSOM* or *Phenograph* algorithms (these algorithms were selected as they are both high performing unsupervised clustering algorithms, an in-depth comparison has previously been described by Weber and Robinson, 2016 [25]). **3)** Data is visualized, and metadata, clusters and cell type labels are used to explore differences between samples/conditions.*A detailed step-by-step guide for using the *ImmunoCluster* tool is available: https://github.com/kordastilab/ImmunoCluster.

## Case studies

This section aims to convey the scope of *ImmunoCluster*, its versatility, and applicability to complex datasets across a range of high dimensional approaches. We implemented the *ImmunoCluster* framework for three different types of high dimensional single cell data (LMC, IMC, and FC). We firstly used previously published and publicly available LMC data [7]. This dataset was chosen as it is representative of an intricate immunophenotyping project and would test the ability of *ImmunoCluster* to reproduce published results. Secondly, we investigated two novel IMC datasets, from HNSCC and DLBCL patients with the aim of demonstrating the applicability of the *ImmunoCluster* tool for IMC data analysis and cell cluster identification and visualization. Finally, FC data from the BM of 7 HDs during hip surgery were used to show the ability of the framework to analyze conventional FC data and identify rare immune cell populations.

### Liquid mass cytometry

The chosen dataset [7] includes mass cytometry data from 15 patients with leukemia who all received BMT, three of these patients developed acute GvHD. Samples were taken at 30 and 90 days post-BMT. Thirty-three cell markers, both lineage and functional were analyzed using mass cytometry (Helios™ CyTOF^®^ system). Prior to uploading into the *ImmunoCluster* framework the FCS files were normalized and gated as described in the methods section.

An initial visualization of the data was first carried out using MDS plots, these were overlaid with metadata to identify the similarity of patients by condition and day of measurement (Figure 3A and 3B). A heatmap was also created to give an overview of marker expression for each patient and annotated with the metadata provided (Figure 3C). The *UMAP* algorithm was used for dimensionality reduction of data. There was a difference in the distribution of cells from GvHD and non-GvHD patients (‘none’) across the cell islands between both timepoints, 30 and 90 days after BMT (Figure 4A).

**Figure 3.**
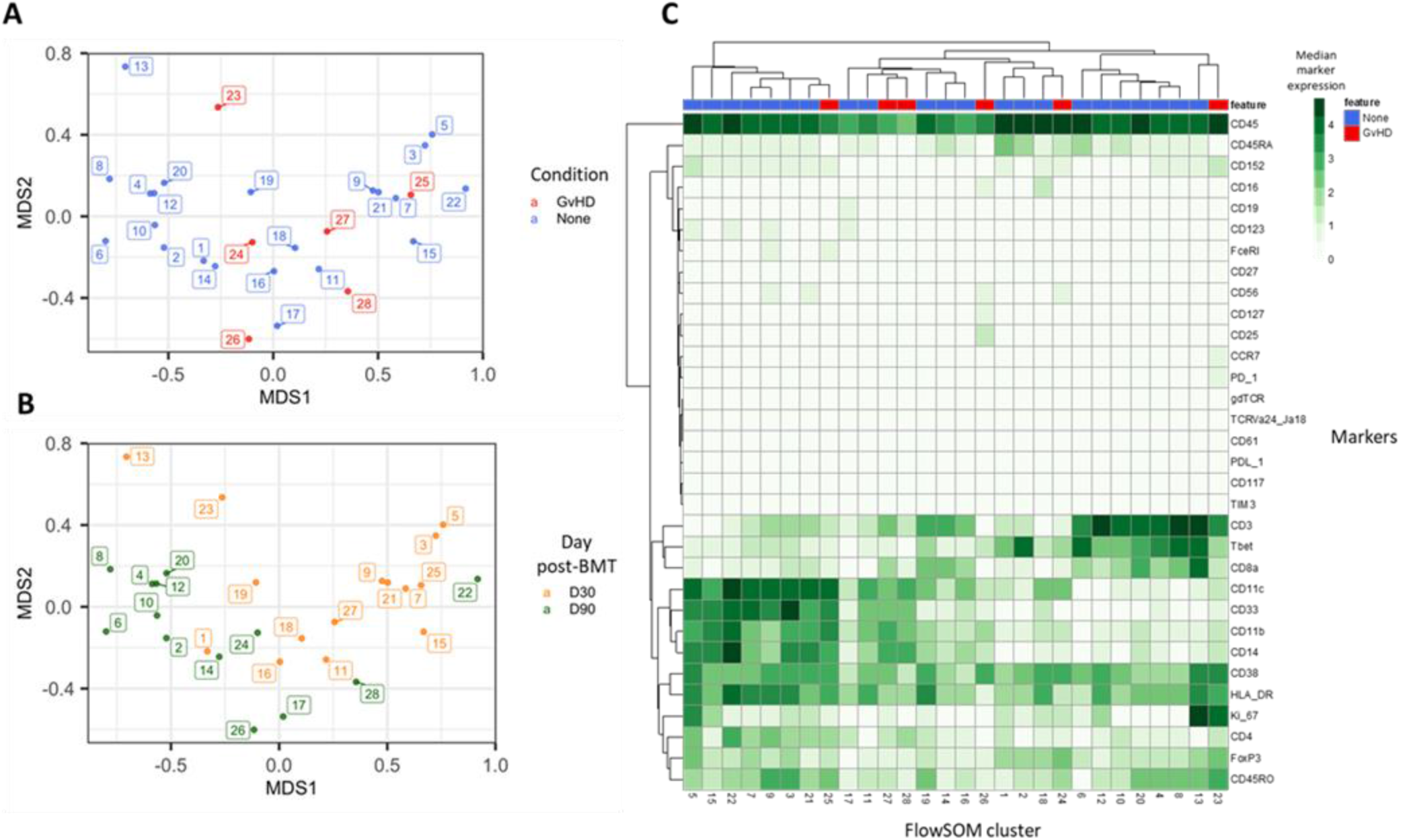
Initial exploration of LMC data from patients with leukemia who received bone marrow transplants. Multidimensional scaling of data, annotated with **A)** condition (GvHD or none), and **B)** time after bone marrow transplant (BMT) treatment. **C)** Heatmap showing the marker expression for each patient.

**Figure 4.**
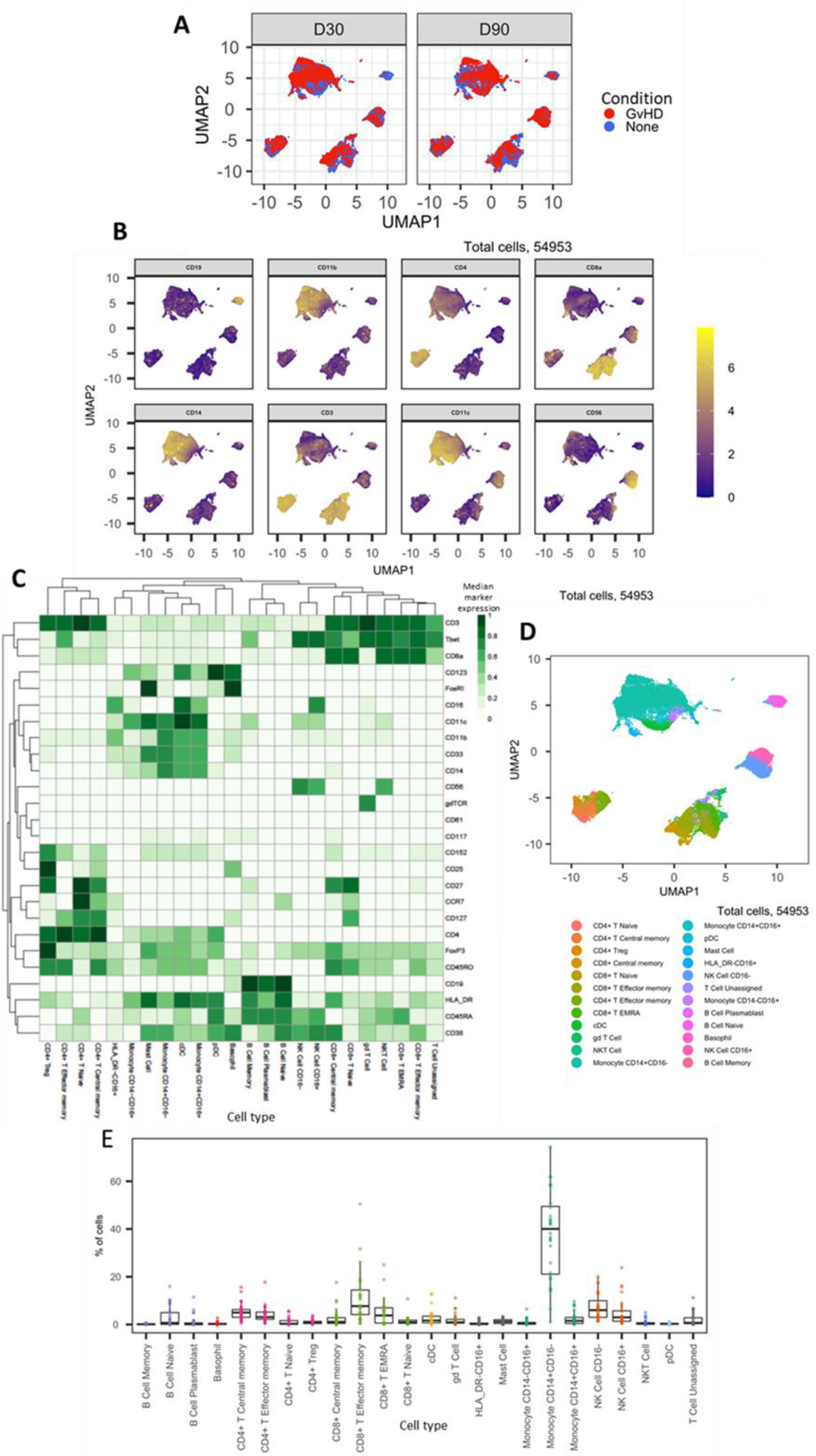
Dimensionality reduced LMC CyTOF data and marker expression: **A)** *UMAP* plots colored by GvHD and none and split by time point (30 and 90 days after BMT treatment). **B)** Expression of eight selected lineage markers projected onto *UMAP* plot. Identifying cell types and abundance of clusters: **C)** Heatmap showing median marker expression across all identified cell types. **D)** *UMAP* annotated with cell types. **E)** Distribution of immune cell frequencies, abundance (%) of each cell type across all samples measured.

The CyTOF^®^ panel used in this experiment was specifically designed to identify all major human immune cell lineages [7], the marker expression across cell islands was visualized and indicated cell types present within the cell islands (Figure 4B) (expression of all markers measured Figure S8). To further distinguish the cell types that were present in the cell islands and visible from the dimensionality reduced *UMAP* plots (Figure 4B) the *FlowSOM* clustering algorithm was applied. The number of clusters input into the *FlowSOM* algorithm was slightly higher than the number of expected clusters (n=56) (Figure S7). The SingleCellExperiment object stored the data for each *K* cluster (*K1*-*K56*), therefore further downstream data exploration was carried out looking at different numbers of *K* clusters. Outputs created showing the expression of the 33 markers measured were used to identify cluster cell types (Figure S7), *ImmunoCluster* was successfully able to replicate the findings from Hartmann *et al.* [7], with 24 of the cell populations being identified (Figure 4C and D). Selecting the correct number of clusters to input into the clustering algorithm can be a challenging aspect of these analyses, but the framework provides the user with multiple parameters to aid this decision. We recommend that users should always overestimate the number of cell populations as data for each *K* number of clusters will be stored in the SingleCellExperiment object and can therefore be explored and refined later using the different visualization techniques available within the framework. Additionally, an elbow plot can be created alongside the clustering algorithm output to guide the decision on the best number of clusters for downstream analysis (Figure S6). Over-clustering can allow for the identification of rare cell types within a dataset, at the expense of often generating several clusters of highly prevalent cell types that likely represent the same biological cell type. *ImmunoCluster* provides the tools to manually and reproducibly merge clusters of the same biological cell identity into one group after over-clustering. These clusters can additionally be merged into each other to use a less granular, higher-level population annotation (see Figure S9 for an example of higher-level clustering).

The abundance of all cell types across all samples was measured, and CD14^+^ CD16^−^ monocytes were identified as the most abundant population, correlating with the Hartmann *et al.* data [7] (Figure 4E). Individually displaying patient’s cell type abundance means that we could identify variation within a group of interest, for example we can see that although there is variation within the GvHD and none, overall, they appear to follow the same trends in cell type abundance (Figure 5A). Additionally, significant differences between memory B-cells (FDR *p*=4.38 × 10^−3^), naïve B-cells (FDR *p*=1.35 × 10^−2^), and naïve CD4^+^ T-cells (FDR *p*=3.47 × 10^−2^) were identified between the GvHD and none (Figure 5B and C). A difference in number of naïve B-cells was previously reported by Hartmann *et al.* [7] in this dataset and can be seen in Figure 4A (using Figure 4D for cell island identification), where there is a noticeable reduction of these cells in the GvHD patients. In addition, a reduction in naïve CD4^+^ T-cells can also be seen, which was also previously reported, but to a lesser extent. A volcano plot can be used to highlight the differentially abundant cell clusters between GvHD and none (GvHD logFC+ve and none LogFC-ve), where cell types with FDR *p*< 0.05 are shown in red (Figure 5D), in line with the previously published data [7]. Additionally, we explored the expression of checkpoint-related molecules, and their receptors, markers of proliferation, and invariant natural killer T (iNK T) cells in CD8^+^ T-cells and compared the expression of these in the patients with GvHD and none (Figure 5E). These markers help assess the functional states of cells, and the checkpoint-related molecules and proliferative activity markers such as PD-1, PD-L1, TIM3, Ki-67, and TCRVa24-Ja18; the expression of these antigens have been proposed previously as candidate biomarkers for immunotherapy [7]. Visually, GvHD patients had a higher expression of PD-1 and a noticeable difference in the Ki-67 proliferation marker, as expected for these patients (Figure 5E).

**Figure 5.**
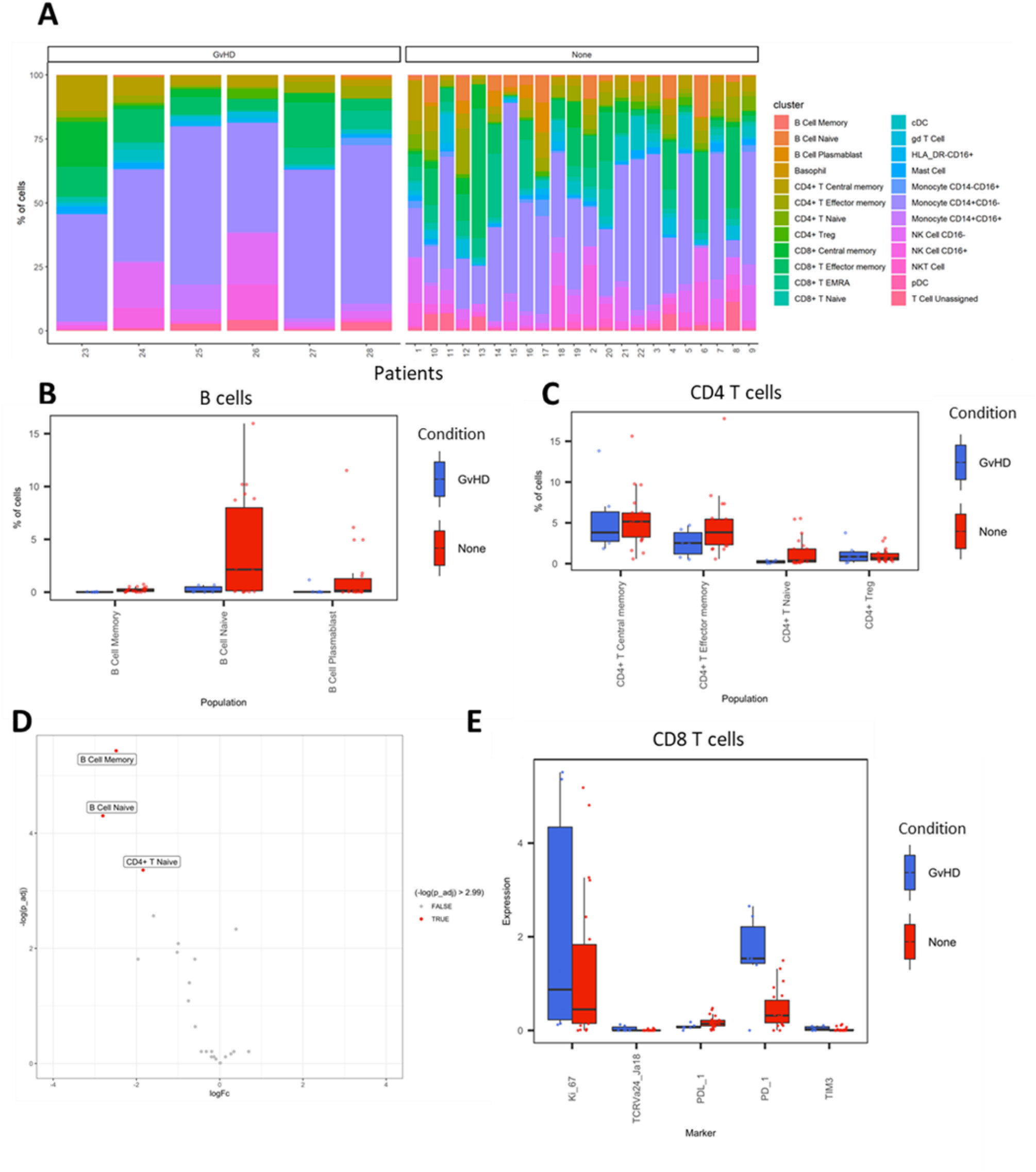
LMC CyTOF data cluster and cell type abundances. **A)** Percentage of each cell type shown for each patient numbered across the bottom of the plot. **B** and **C)** Boxplots portray IQR with the horizontal line representing the median percentage of cell types in both GvHD and non-GvHD patients for B-cell and CD4^+^ T-cell populations, respectively. **D)** Differential abundance analysis; volcano plot showing the significantly differentially expressed cell abundances (FDR P < 0.05) between GvHD and none (GVHD logFC+ve and none LogFC-ve). **E)** Comparison of checkpoint-related molecules (PD-1, and TIM3), receptors (PD-L1), proliferative (Ki-67), and iNK T-cells (TCRVa24-Ja18) marker expression between GvHD and non-GvHD (none) patients in the CD8^+^ T-cell cluster.

### Imaging mass cytometry datasets

We utilized two different IMC datasets to demonstrate how *ImmunoCluster* would handle IMC data from two different tumor microenvironments which possess, a) a biological image with clear boundaries, such as HNSCC, where clear tumor and stroma regions are evident; b) a biological image which was more heterogeneous and diffuse, such as the DLBCL lymph node section. *ImmunoCluster*’s ability to explore IMC data adds a novel element to the framework and demonstrates the flexibility and applicability of the tool. This unique element within the framework means that researchers can explore IMC data easily, in-line with LMC and FC data. After data pre-processing (see methods and Figure S2), the transformed gated FCS files were uploaded into the *ImmunoCluster* framework. Generally, the same workflow which was applied to the LMC and FC data was applied to the IMC data from a section of tissue from patients with HNSCC and DLBCL (lymph node), as a proof of principle application of the *ImmunoCluster* tool for IMC data (Figures 6 and 7). The only difference with the IMC data was the use of a rank heatmap, this ranks the order of each marker for each sample, (1-total number of samples). The reason for this was for clarity and ease of cell identification across clusters as IMC produces lower resolution data compared to the data output of LMC or FC, this makes manual assignments of populations more difficult to interpret. Therefore, a rank of expression was used as an easy means to define the highest and lowest expression of markers between clusters.

**Figure 6.**
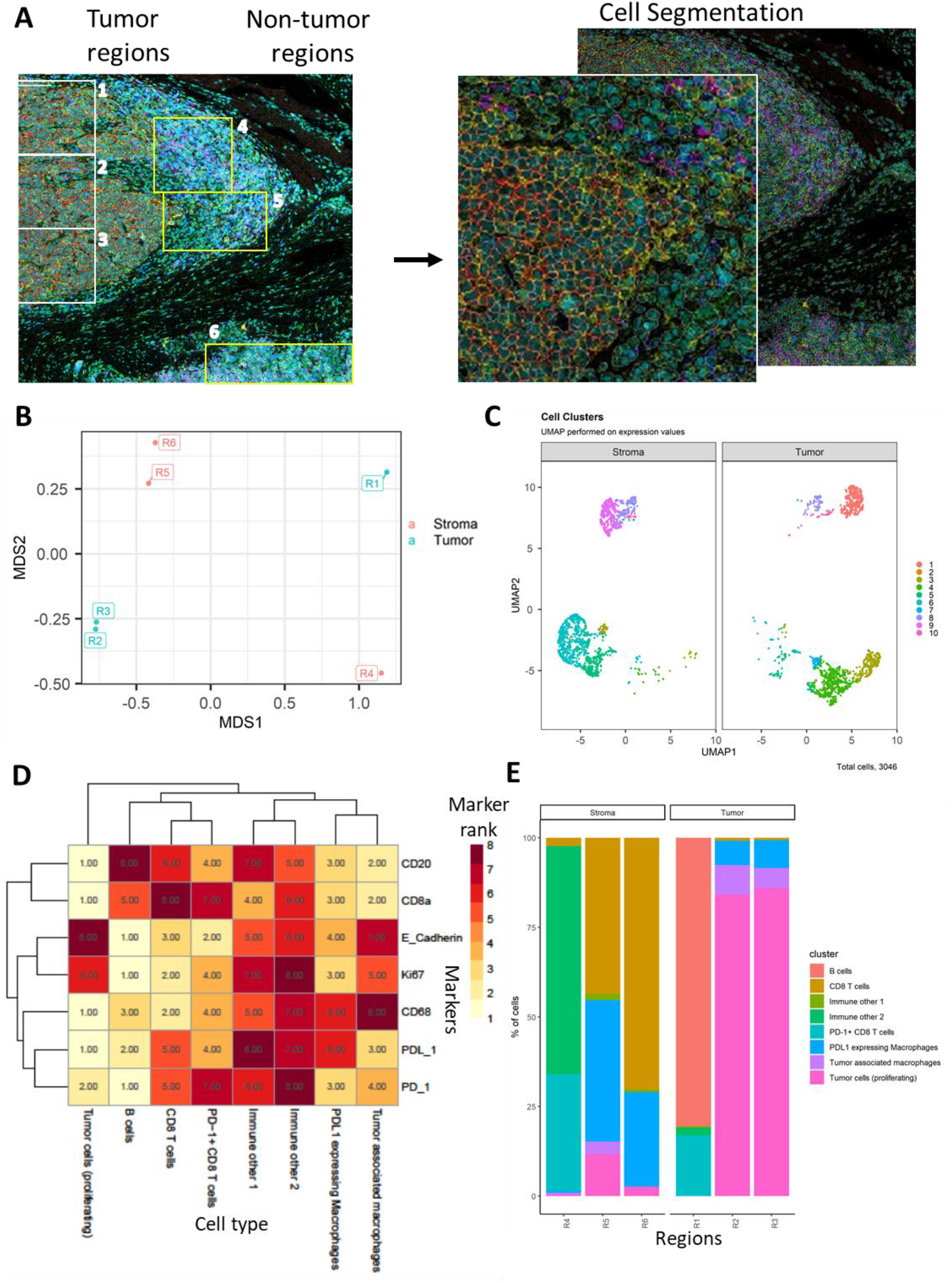
Head and neck IMC data: Immunophenotyping the tumor microenvironment with IMC data using *ImmunoCluster*. **A)** IMC image showing an example 5 channels: PD-L1 (green), CD4 (yellow), E-Cadherin (red), CD20 (magenta) and CD8α (blue). The tumor and stroma areas are clear to the eye and 3 regions were selected from each as shown (regions 1-6). Images with the segmented cell borders highlighted. **B)** MDS plot of stroma and tumor regions. **C)** Dimensionality reduced data (*UMAP* algorithm applied) annotated with *FlowSOM* clusters and split by region type. **D)** Rank heatmap: ranked expression (1-8, where 8 is high) of 9 markers (CD20, CD8α, CD4, E-Cadherin, Ki-67, PD-L1, CD68, and PD-1) and identified cell type. **E)** Proportion of cell types for each tissue region.

**Figure 7.**
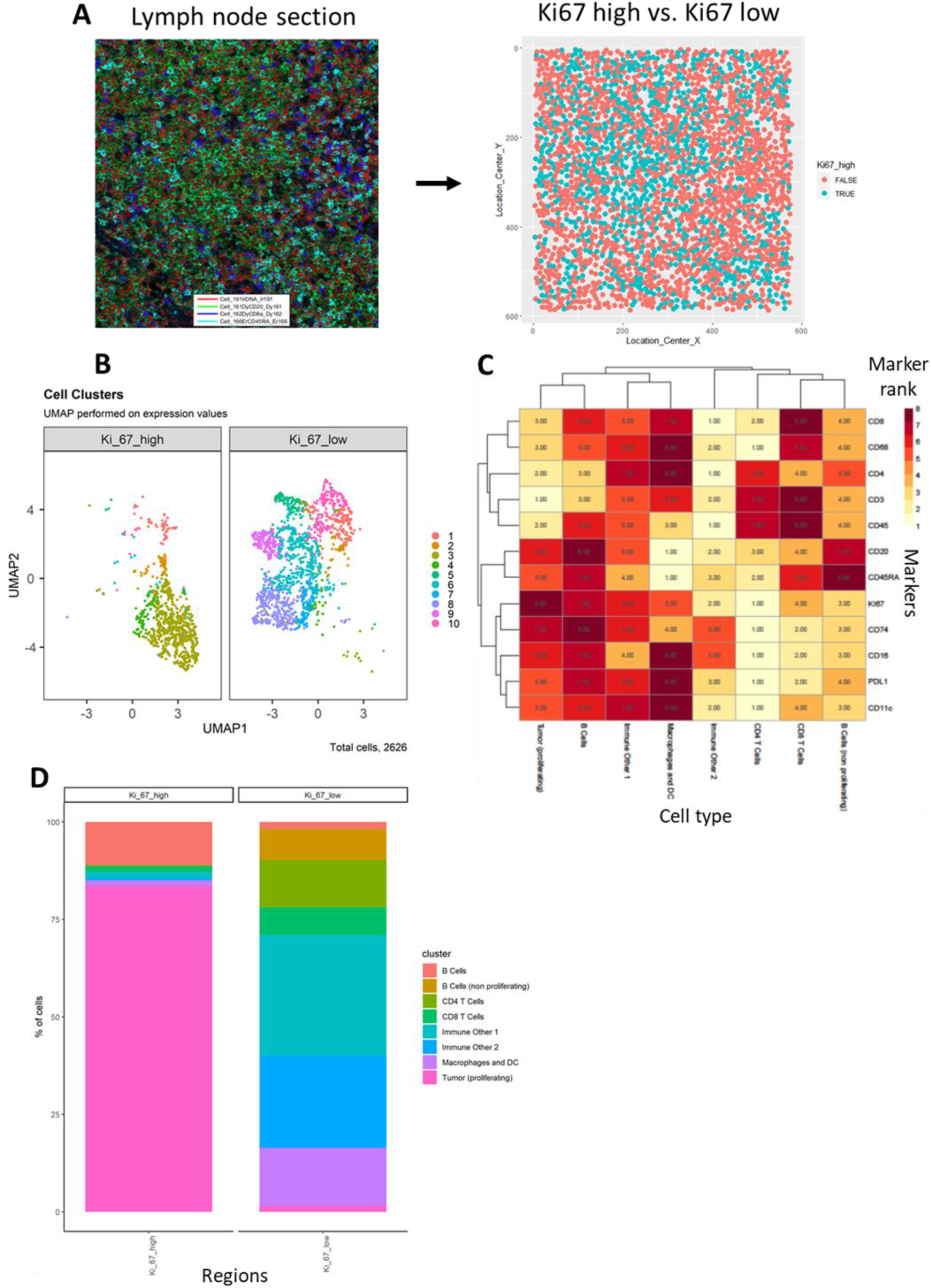
Diffuse large B-cell lymphoma IMC data: Immunophenotyping the lymph node microenvironment with IMC data using *ImmunoCluster*. **A)** IMC image showing an example 4 channels: DNA (red), CD20 (green), CD8α (dark blue), and CD45RA (light blue). The image was split by high and low Ki-67 expression to identify highly proliferative cells (tumor). **B)** Dimensionality reduced data (*UMAP* algorithm applied) annotated with *FlowSOM* clusters and split by Ki-67 high or low. **C)** Rank heatmap: ranked expression (1-8, where 8 is high) of 12 markers (CD8, CD68, CD4, CD3, CD45, CD20, CD45RA, Ki-67, CD74, CD16, PD-L1, and CD11c) and identified cell type. **E)** Proportion of cell types for Ki-67 high and Ki-67 low cell populations.

#### Head and neck squamous cell carcinoma tissue section

Cell segmented regions were annotated in Figure 6A, regions 1-3 were selected as tumor regions and 3-6 as stroma regions. The *ImmunoCluster* tool was able to distinguish between the stroma and tumor regions of the tissue section from the HNSCC tumor (Figure 6B), and with additional markers could provide more in-depth analysis of the cell phenotypes in each respective region of the tumor. Dimensionality reduction of the data clearly separated the cell islands belonging to the stroma and tumor regions of the tissue (Figure 6C), suggesting different cell phenotypes between the regions. The *FlowSOM* algorithm was applied to cluster the cells and identify cell populations. The rank heatmap was used to show the ranked expression of each marker (ranked 1-10, with 10 being high) for each cluster (1-10) (Figure S10), identified cell types were then used to annotate the heatmap (Figure 6D). High expression of E-cadherin can be used as a marker of cancerous tissue in HNSCC, and its expression correlates with the cancer regions selected for analysis (Figure 6D). We focused on CD8^+^ T-cell, B-cell, and macrophage cell clusters [26, 27]. Regions 2 and 3 of the tumor mostly consist of proliferating (Ki-67^+^) tumor cells and macrophages (tumor-associated and PD-L1^+^), Region 1 was a mix of immune cells, B-cells, and PD-L1^+^ CD8^+^ T-cells (Figure 6E). Stromal regions 5 and 6 are mostly PD-L1^+^ macrophages and CD8^+^ T-cells. Region 4 has a cluster of mixed immune cells, and from the IMC image (Figure 6A) we can see this region looks like the tumor is encroaching into the stromal region. The stroma seems to be mostly non-proliferating cells (KI-67^−^ compared to the tumor region) and CD8^+^ T-cells, including a CD20^+^ CD8^+^ cytotoxic subset [28].

#### Diffuse large B-cell lymphoma lymph node section

The IMC lymph node section was split by high and low Ki-67 expression to identify the highly proliferative tumor cells (Figure 7A); Ki-67 is used as a prognostic marker in DLBCL [29]. The *FlowSOM* algorithm was applied to the data to split the IMC data into clusters, these clusters were projected onto the *UMAP* plot to visualize how these clusters were split between the Ki-67 high and low (Figure 7B). The marker expression ranking tool was applied to the clustered data and used to identify cell types (Figure S11 and 8C). The majority of Ki-67 high population were proliferating tumor cells (84%), and the Ki-67 low population consisted of a heterogeneous collection of immune cell populations, such as CD4^+^ T cells, CD8^+^ T-cells, B-cells, macrophages and dendritic cells (DCs) that were successfully identified using the *ImmunoCluster* tool (Figure 7D).

### Flow cytometry

The healthy donor BM taken during hip surgery was gated for the CD3^+^ CD4^+^ population before analysis with in the *ImmunoCluster* framework, with the aim of testing *ImmunoCluster’s* ability to identify minor populations such as Tregs and their subpopulations [30]. The *FlowSOM* algorithm was applied to the data resulting in 40 clusters (Figure S12), and a heatmap was created with median expression of markers to identify cell types (Figure 8B). Due to the markers used and the low prevalence of Tregs the majority of cells were identified as CD4^+^ T-cells. Additionally, three populations of Tregs were identified by the *ImmunoCluster* tool; Tregs (CD25^+^ CD127^low^) (3.6%, 1.3-5.5), Treg A (CD25^+^, CD127^low^, CD45RA^+^) (0.7%, 0.1-2.0) and Treg B cells (CD25^+^, CD127^low^, CCR4^+^, CD95^high^) (0.9%, 0.2-2.4) (Figure 8C). The abundance of Treg A and B for all healthy donors is shown in Figure 8D and their distribution was as expected [30].

**Figure 8.**
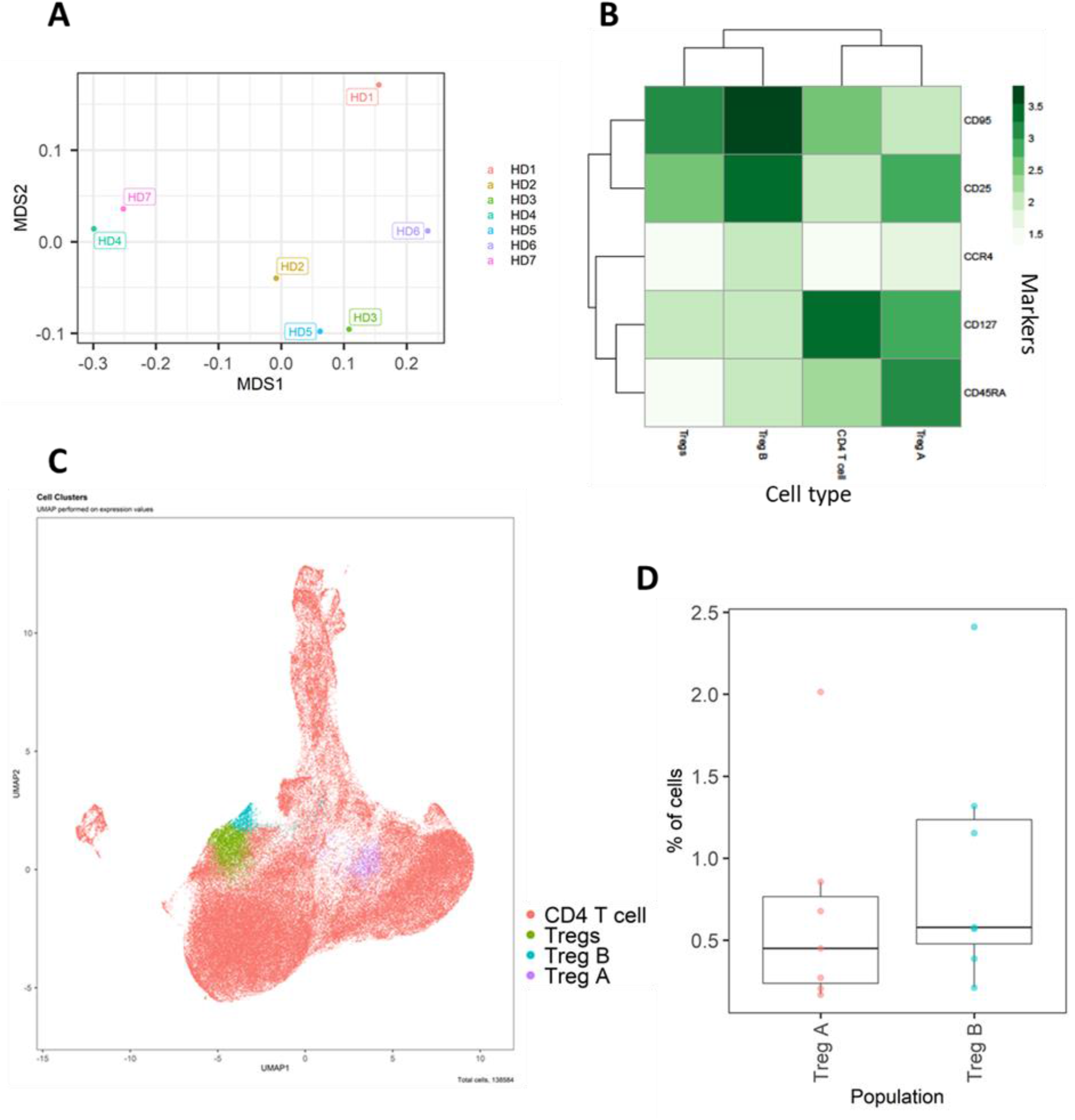
Healthy donor bone marrow flow cytometry data: identification of rare CD4^+^ T-cell immune cell subsets. **A)** MDS plot of each healthy donor (1-7). **B)** Heatmap showing marker expression across clusters of identified cell types. **C)** Dimensionality reduced data (*UMAP* algorithm applied) annotated with cell type. **D)** Proportion of Treg A and B from total cells from healthy donors.

## Discussion

Here we provide an overview of the flexibility and scope for utilizing *ImmunoCluster*, an open-access non-specialist framework using LMC, IMC and FC datasets. We have emphasized the ease-of-use, clinical utility, and quality of outputs which can be created. Explanations of the functions and workflow have been provided and an in-depth step-by-step protocol and tutorial are available on the GitHub site (https://github.com/kordastilab/ImmunoCluster).

The purpose for creating the *ImmunoCluster* framework was to support researchers carrying out complex immunophenotyping experiments. The dawn of ‘big data’ and its ever-growing utilization has surpassed the number of experimental parameters that a researcher can feasibly analyze as a collective. As computational training is still a niche area, the *ImmunoCluster* framework was designed in collaboration between wet lab and computational biology scientists, with the purpose of creating a tool which was sophisticated in its analysis, yet easy to use in its application. We provide sufficient detail and explanation for the researcher to understand the need for each analysis step as well as how to confidently execute the process.

*ImmunoCluster* has been built with an emphasis on generalizability and scalability to facilitate a broad use and is an advance over other open-source tools as it provides an integrated framework to uniquely perform a complete computational pipeline on high-dimensional LMC, IMC and FC data. The inclusion of primary data analysis for IMC brings a unique element to the current protocol, which in addition to its highly developed ease-of-use interface, differentiates it even further from other published protocols, including data QC, exploration, analysis and visualization; with an aim of performing transparent and reproducible analyses. *ImmunoCluster* was built to easily leverage many popular R packages for flow cytometry data (i.e. *RPhenograph*, *Rtsne* and *FlowSOM*), selected because they are open-source and regularly maintained with extensive documentation. All methods prioritize simple, understandable and attractive visualizations designed to be used by both dry- and wet-lab researchers/clinicians. The computational methods used within the framework are importantly readily scalable to a dataset of millions of cells across several samples and a variety of experimental or phenotypic conditions. *ImmunoCluster* can also be used on a local desktop with standard configuration, and depending on the number of cells, can usually be run within a day end-to-end.

The limitations of this computational framework are important to take into consideration when designing an experiment. Unsupervised Clustering (Stage 2) of samples is the most important stage of the framework and its ability to accurately define populations across all samples is critical to all later stages of investigation. The number of *k* clusters to generate (cluster resolution) can significantly affect the ability to identify or determine a change in a biologically relevant population. Merging two subpopulations due to low resolution of clustering may mask an important experimental observation. Determining the precise number of clusters that are relevant for a given dataset is an important step. In our step-by-step guide we provide complete documentation and clear user-friendly approaches to optimally define this.

Technical inter-sample marker signal variability due to batch effects may impact the ability of unsupervised analysis to reliably detect certain populations, if significant batch effects are present. One approach commonly employed in mass cytometry to overcome this is to use sample barcoding, reducing the variability between each sample, allowing all samples to be exposed to the same antibody mixture [31]. If there are significant differences in the total number of cells recovered between samples, samples with many more cells may bias the clustering. Samples with very few cells recovered may result in information loss and missing populations that are in fact present. Although channel spillover is diminished in mass cytometry, it still exists in fluorescence flow cytometry, and should be considered when designing antibody panels to reduce the effects on introducing cell phenotype artifacts in downstream unsupervised analysis. As such, initial exploratory data analysis is key to determining if any of these confounding factors might be present in the data.

## Summary

The *ImmunoCluster* package increases the accessibility of advanced computational methods for users tasked with generating high-quality analysis of high-dimensional LMC, IMC, and FC data. More advanced users can leverage *ImmunoCluster*’s data structures, methods and suitability for scripted analysis to integrate our work with other R analysis tools and build on the package’s foundation. In our clinically relevant immune monitoring case study setting, the *ImmunoCluster* framework successfully identified 24 phenotypically distinct cell clusters, and their abundance across all samples, and highlighted differentially represented cells between GvHD and non-GvHD patients, all of which were consistent with the Hartmann *et al.* data [7]. We also applied the framework to two sets of IMC data, showing the broad applicability of *ImmunoCluster* as well as the ease of being able to analyze the IMC data in-line with LMC data, helping compare these types of data, which will be useful in future studies where multiple technologies are being applied within one study. Finally, we implemented FC data into *ImmunoCluster*, confirming that it was able to analyze conventional FC data and identify rare immune cell populations. Currently the pre-processing of raw files is carried out prior to implementation into the *ImmunoCluster* framework, but due to the flexibility and design of the tool, future implementations may include IMC data pre-processing within the *ImmunoCluster* R package. This versatile framework provides an opportunity to include all researchers with varying knowledge and experience in computational biology to be involved in the experimental project from experimental design, wet lab/clinical trial, all the way through the data analysis process and visualization.

