## Supplementary for "ImmunoCluster: A computational framework for the non-specialist to profile cellular heterogeneity in cytometry datasets"

**Table S1.** Reference panel of anti-human antibodies for mass cytometry used by Hartmann *et al.* [7].

| Isotope | Element | Marker | Clone | Staining step |
|---|---|---|---|---|
| 89 | Y | CD45 | H130 | Surface |
| 139 | La | CD235αβ/CD61 | HIR2/VI-PL2 | Surface |
| 141 | Pr | CD3 | UCHT1 | Surface |
| 142 | Nd | CD19 | HIB19 | Surface |
| 143 | Nd | CD117 | 104D2 | Surface |
| 144 | Nd | CD11b | IRCF44 | Surface |
| 145 | Nd | CD4 | RPA-T4 | Surface |
| 146 | Nd | CD8α | RPA-T8 | Surface |
| 147 | Sm | CD11c | BU15 | Surface |
| 148 | Nd | CD14 | RMO52 | Surface |
| 150 | Nd | FcεRI | AER-37 (CRA-1) | Surface |
| 151 | Eu | CD123 | 6H6 | Surface |
| 152 | Sm | γδTCR | 11F2 | Surface |
| 153 | Eu | CD45RA | HI100 | Surface |
| 154 | Sm | TIM3 | F38-2E2 | Surface |
| 156 | Gd | PD-L1 (CD274) | 29E.2A3 | Surface |
| 158 | Gd | CD27 | L128 | Surface |
| 160 | Gd | Tbet | 4B10 | Intracellular |
| 161 | Dy | CD152 (CTLA-4) | 14D3 | Intracellular |
| 162 | Dy | FoxP3 | PCH101 | Intracellular |
| 163 | Dy | CD33 | WM53 | Surface |
| 164 | Dy | CD45RO | UCHL1 | Surface |
| 165 | Ho | CD127 | A019D5 | Surface |
| 167 | Er | CCR7 (CD197) | G043H7 | Surface |
| 168 | Er | Ki-67 | B56 | Intracellular |
| 169 | Tm | CD25 | 2A3 | Surface |
| 170 | Er | TCR Va24-Ja18 | 6B11 | Intracellular |
| 172 | Yb | CD38 | HIT2 | Surface |
| 174 | Yb | HLA-DR | L243 | Surface |
| 175 | Lu | PD-1 | EH12.2H7 | Surface |
| 176 | Yb | CD56 | NCAM16.2 | Surface |
| 209 | Bi | CD16 | 3G8 | Surface |

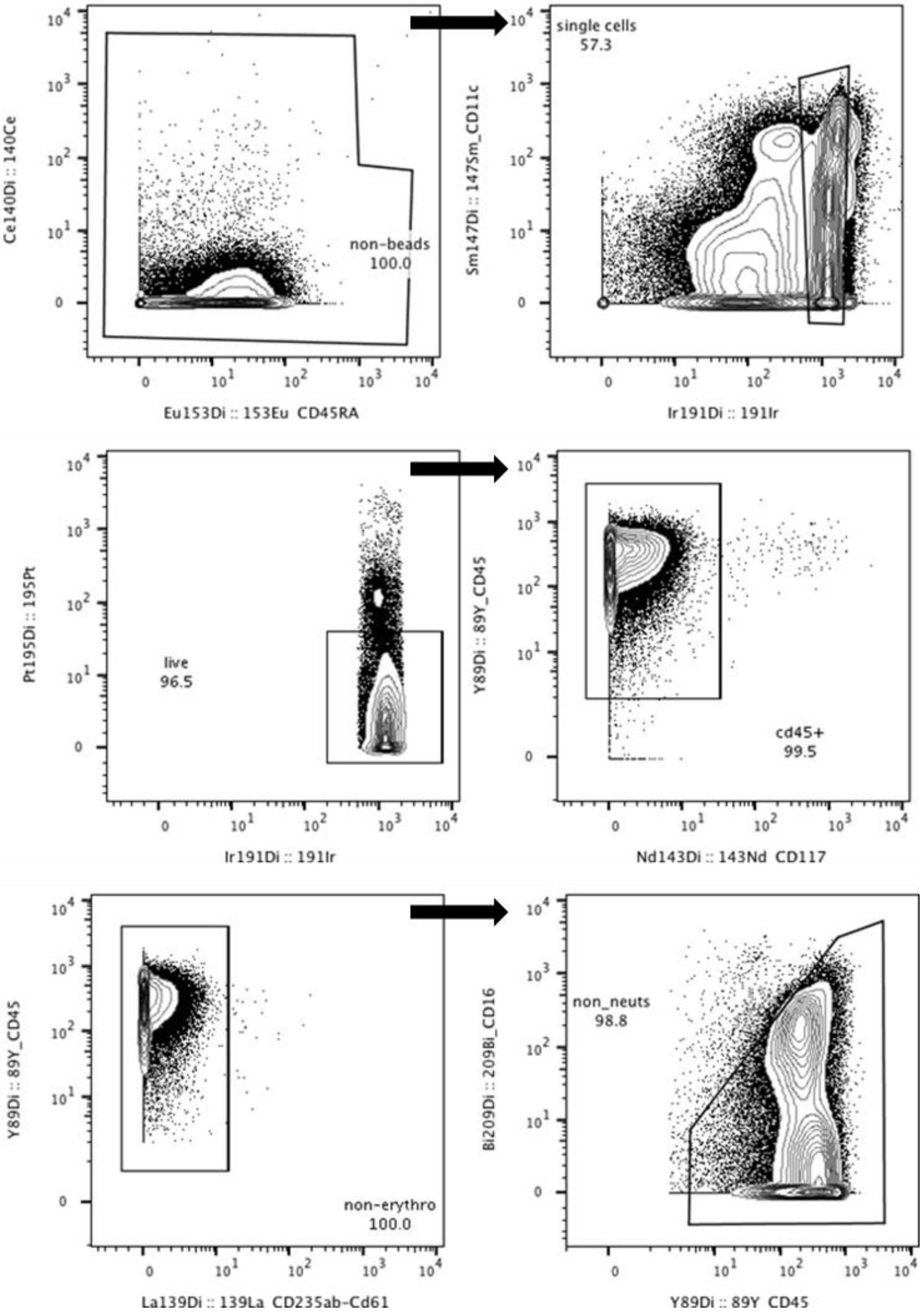

**Figure S1.** Mass cytometry data gating strategy in FlowJo. Calibration beads, doublets, dead cells, non-CD45$^+$ cells, erythrocytes (CD235αβ/CD61$^+$), and neutrophils (CD16$^+$) were removed.

**Table S2.** Reference panel of anti-human antibodies for the head and neck cancer (HNC) imaging mass cytometry experiment.

| Isotope | Element | Channel | Marker |
|---|---|---|---|
| 150 | Nd | C1 | PDL-1 |
| 152 | Sm | C2 | CD45 |
| 156 | Gd | C3 | CD4 |
| 158 | Gd | C4 | E-Cadherin |
| 159 | Tb | C5 | CD68 |
| 161 | Dy | C6 | CD20 |
| 162 | Dy | C7 | CD8α |
| 165 | Ho | C8 | PD-1 |
| 168 | Er | C9 | Ki-67 |
| 170 | Er | C10 | CD3 |
| 191 | Ir | C11 | DNA |
| 193 | Ir | C12 | DNA |

**Table S3.** Reference panel of anti-human antibodies for the diffuse large B-cell lymphoma (DLBCL) imaging mass cytometry experiment.

| Isotope | Element | Marker | Dilution* |
|---|---|---|---|
| 141 | Pr | αSMA | 1:2000 |
| 144 | Nd | CD74 | 1:100 |
| 146 | Nd | CD16 | 1:200 |
| 147 | Sm | CD68 | 1:500 |
| 150 | Nd | PD-L1 | 1:100 |
| 151 | Eu | CD31 | 1:200 |
| 152 | Sm | CD45 | 1:500 |
| 155 | Gd | FOXP3 | 1:1000 |
| 156 | Gd | CD4 | 1:200 |
| 161 | Dy | CD20 | 1:400 |
| 162 | Dy | CD8 | 1:400 |
| 165 | Ho | PD1 | 1:100 |
| 166 | Er | CD45RA | 1:2000 |
| 167 | Er | Granzyme B | 1:100 |
| 168 | Er | Ki-67 | 1:1000 |
| 170 | Er | CD3 | 1:400 |
| 173 | Yb | CD45 RO | 1:1000 |
| 175 | Lu | CD11c | 1:400 |
| 176 | Yb | β2 microglobulin | 1:200 |
| 191 | Ir | nuclei | 1:4000 |

*Dilution factors for antibody cocktail described on File S2.

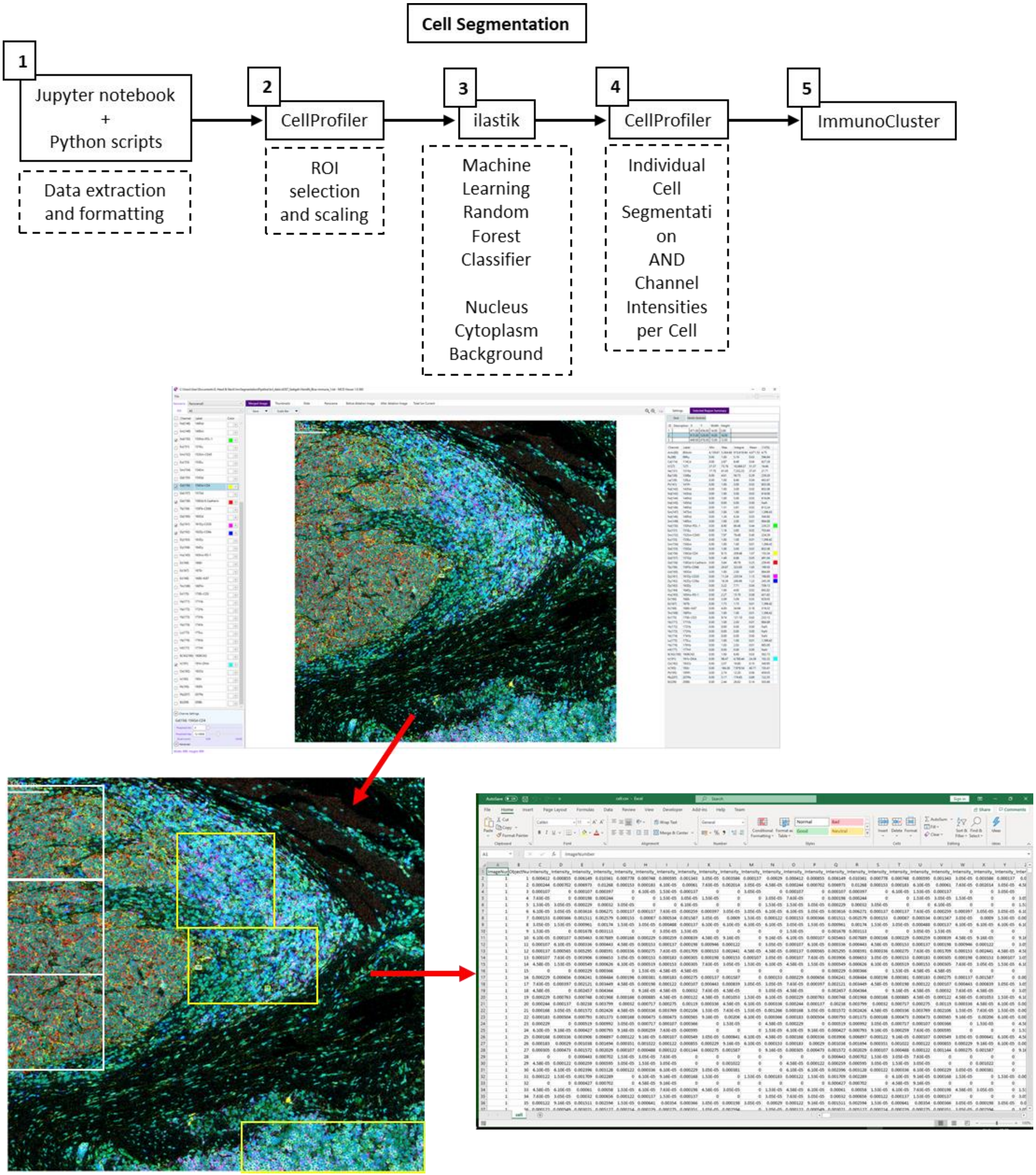

**Figure S2.** IMC data pre-processing work flow. 1) The imctools Python package [11] was used to convert raw IMC files (.mcd, .txt) into intermediatory .tiff files which were used as input files for the following tools. 2) .tiff files were imported into CellProfiler for regions of interest (ROI) selection for classifier training. 3) Ilastik was used for pixel classification, pixels were identified as nuclear, cytoplasmic, or background and these class probabilities were exported as RGB (red, green, blue) .tiff images. 4) The

Ilastik RGB probablilities and the original images were imported into CellProfiler for single-cell segmentation, mask generation and marker quantification.

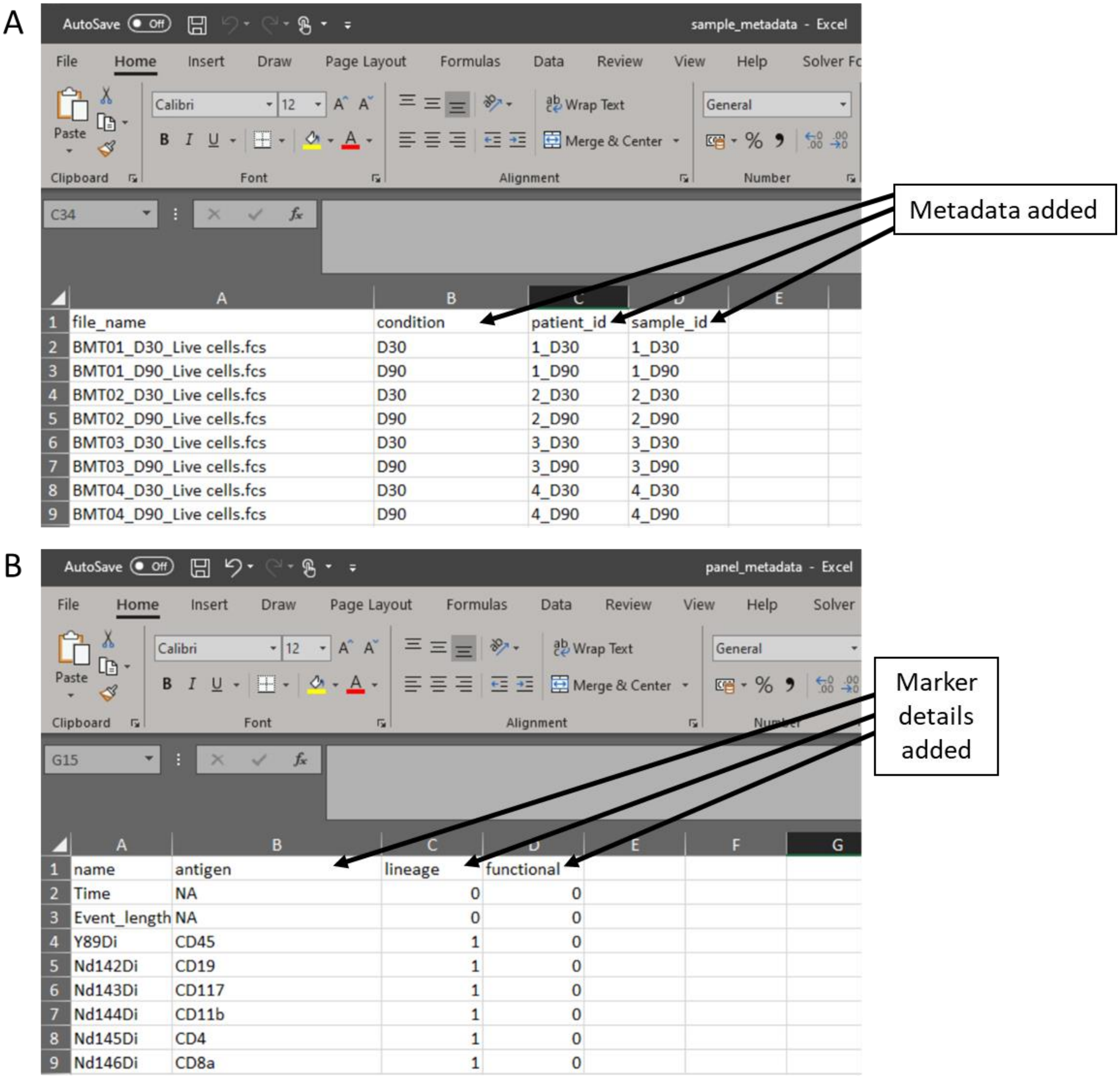

**Figure S3.** Metadata files created in the experimental design stage. A) Sample metadata file containing all metadata the researcher would like to explore throughout the data analysis stage. B) Panel metadata files allows the researcher to re-name parameters and select the markers which will be used for the dimensionality reduction of the data.

### 2. Data Exploration

We provide several functions to examine global patterns within the data, they can be used to get an idea of the influence of conditions/treatment or batch effects within the data. The mdsplot() function will generate a multi-dimensional scaling (MDS) plot on median expresion values for each channel. MDS plots represent an unsupervised way to indicate similarities between samples at a global level berfore more in depth analsysis. In the example here we can see that there is no observable high-level difference between the two GvHD condition samples, however there does appear to be a weak separation between samples collected at D30 and D90 post BMT. Here and throughout the pipeline, colkey can be used to specify plotting colors for graphing conditions.

```
mds1 = mdsplot(sce_gvhd,
               feature = "condition",
               colkey = c(None = 'royalblue', GvHD = 'red2'))

mds2 = mdsplot(sce_gvhd,
               feature = "day_id",
               colkey = c(D30  = 'darkorange1', D90 = 'darkgreen'))

plot_grid(mds1, mds2,
    labels = c('A','B'),
    ncol = 2, align = "l", label_size = 20)
```

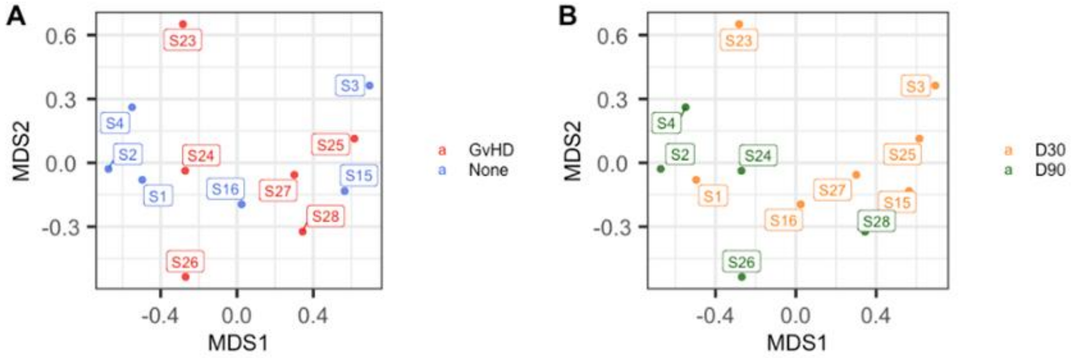

### 3. Dimensionality reduction and clustering

immunoCluster supports multiple different dimensionality reduction algorithms and has UMAP and tSNE functionality implemented below with the ability to downsample before running the algorithms. We run both UMAP and tSNE on selected lineage markers, downsampling to 1000 randomly selected cells per sample to reduce runtime. The dimensionality reductions are stored in the reducedDimNames slot and any other dimensionality reduction, like PCA, can also be stored in the SCE object in parallel.

```
require(umap)
# Run UMAP and store in sce object
sce_gvhd = performUMAP(sce_gvhd, downsample = 1000, useMarkers = clustering_markers)

# Run tSNE and store in sce object
sce_gvhd = performTSNE(sce_gvhd, downsample = 1000, useMarkers = clustering_markers)

sce_gvhd
```

```
## class: SingleCellExperiment
## dim: 32 912813
## metadata(5): file group condition patient_id day_id
## assays(1): scaled
## rownames(32): CD16 CD152 ... HLA_DR CD56
## rowData names(0):
## colnames(912813): cell1 cell2 ... cell912812 cell912813
## colData names(0):
## reducedDimNames(2): UMAP Rtsne
## spikeNames(0):
## altExpNames(0):
```

**Figure S4.** Dimensionality reduction: examples taken from the Github page (https://github.com/kordastilab/ImmunoCluster) highlighting the three different types of dimensionality reduction algorithms available for the users. MDS is part of the initial data exploration step followed by *UMAP* and *tSNE* dimensionality reduction in Stage 2 of the *ImmunoCluster* framework.

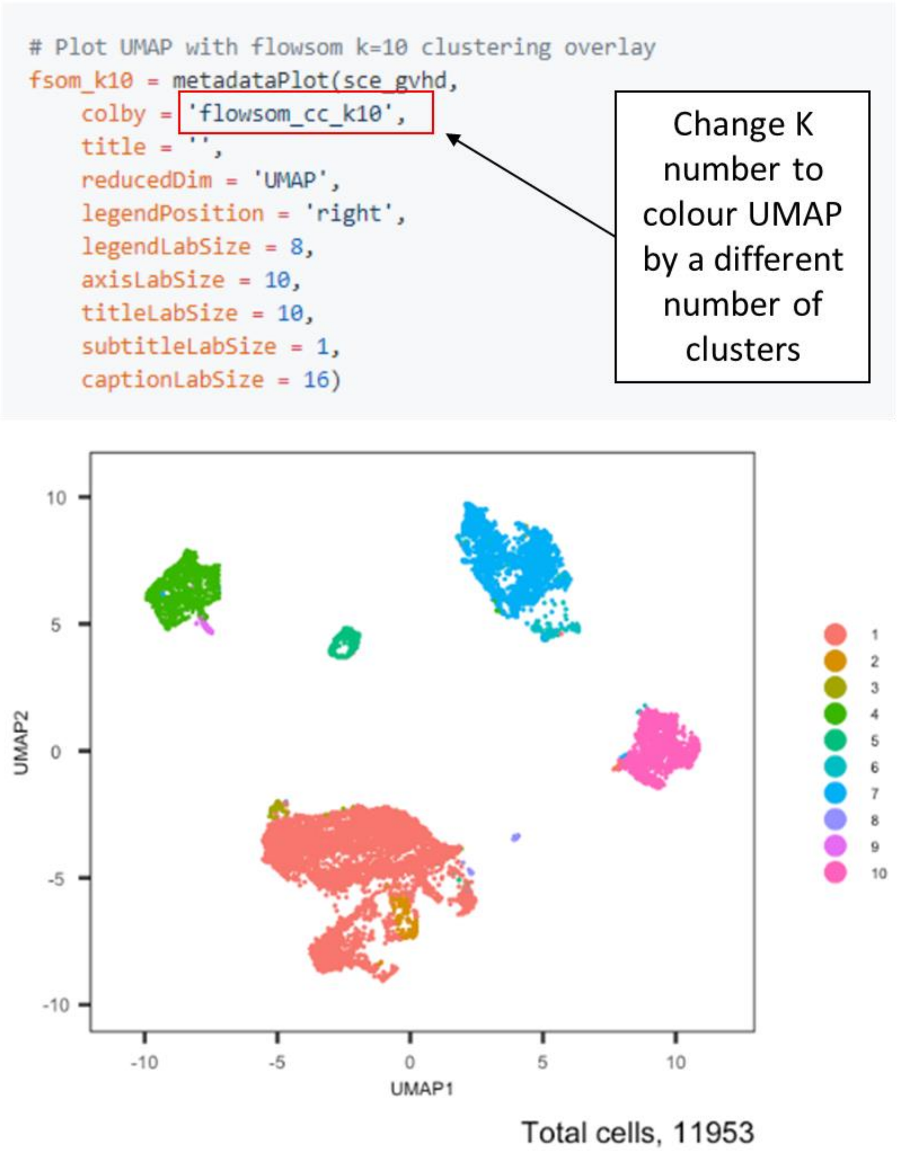

**Figure S5.** Selecting *K* clusters for visualization and downstream anlysis. The *ImmunoCluster* tool saves all *K* clusters selected for the FlowSOM algorithm (e.g. 1-60 *K* clusters), therefore reserachers can view different numbers of *K* clusters for downstream analysis by changing the Flowsom_cc_K number (highlighted above).

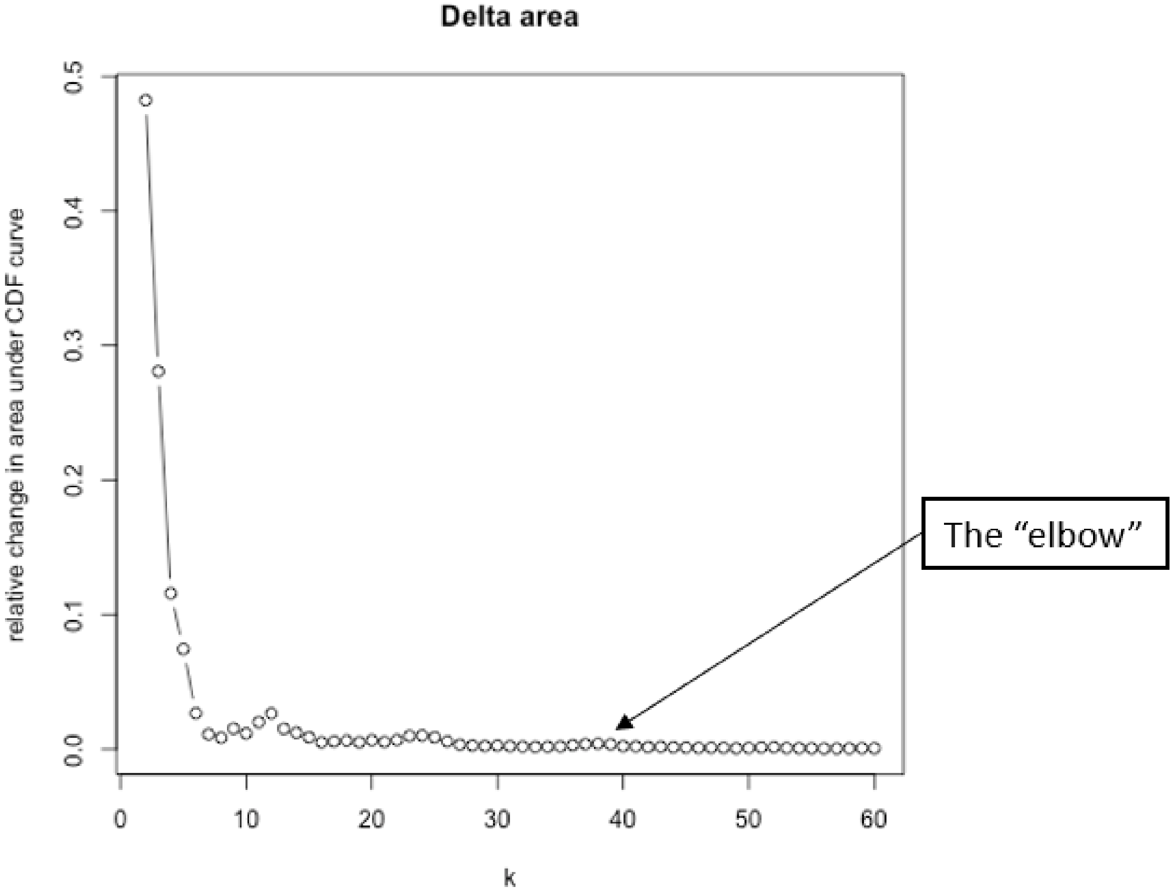

**Figure S6.** Elbow plot criterion to help determine optimal number of clusters for *K*-means clustering (*FlowSOM*). The figure shows *K* clusters 2-60, for each value of *K* the sum of squared errors (SSE) were plotted, the aim was to detect the “elbow”, which is the point where the variance stops decreasing sharply on the plot, representing the best value of *K*.

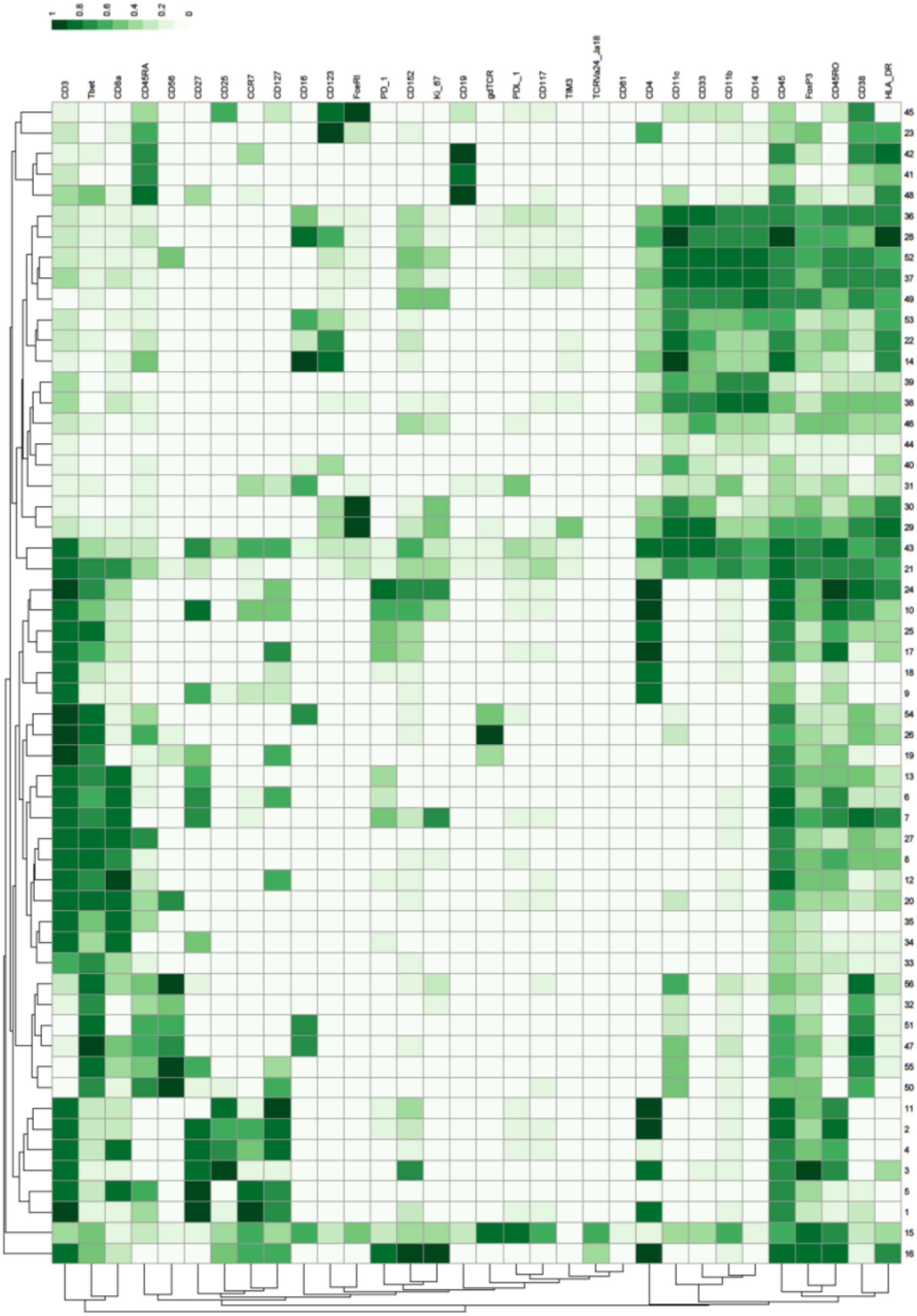

**Figure S7.** Heatmap showing the marker expression of 56 *FlowSOM* clusters. This heatmap was used to identify the cell type of each cluster.

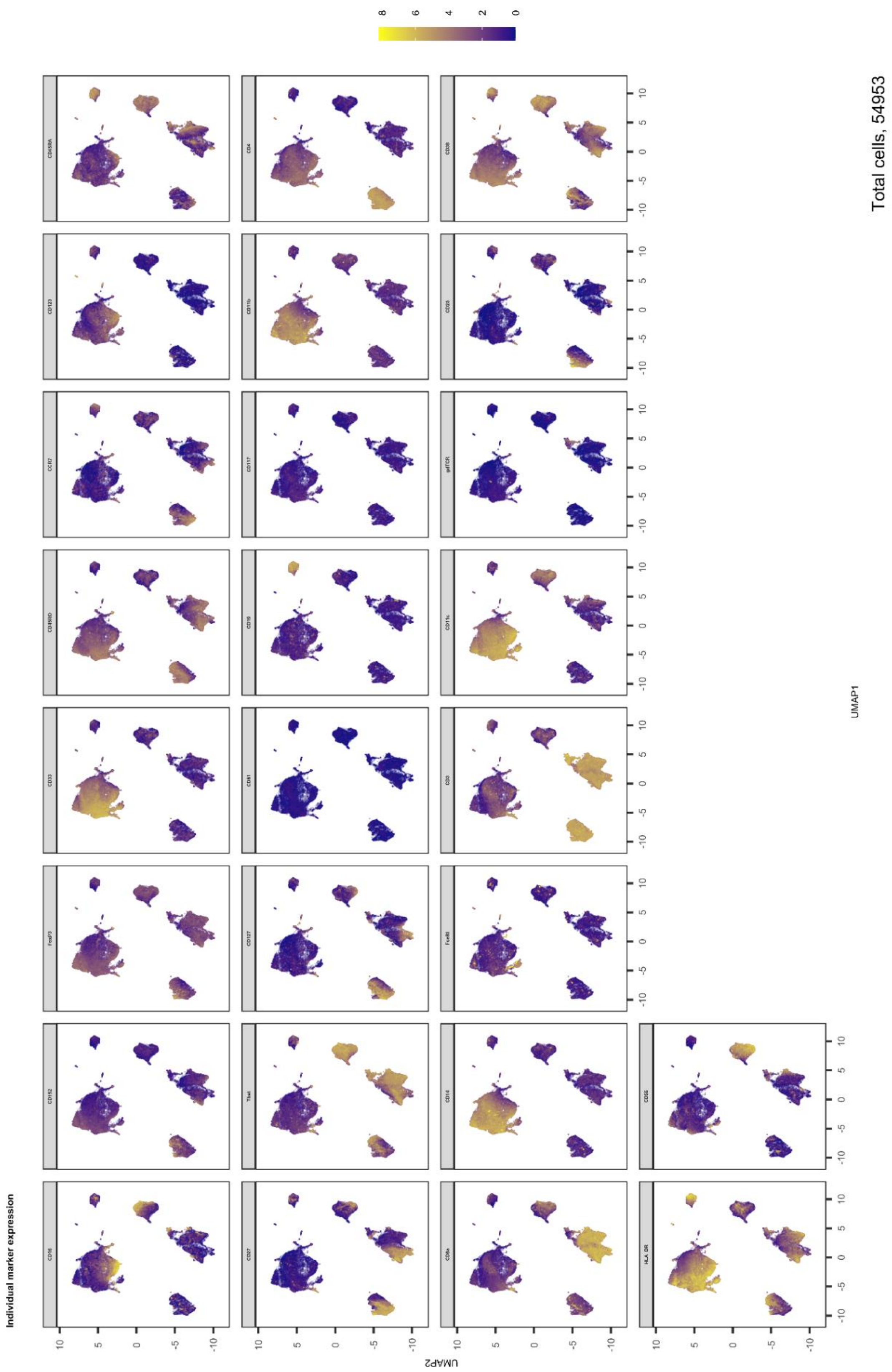

**Figure S8.** Expression of all markers measured projected onto cell islands produced by *UMAP*.

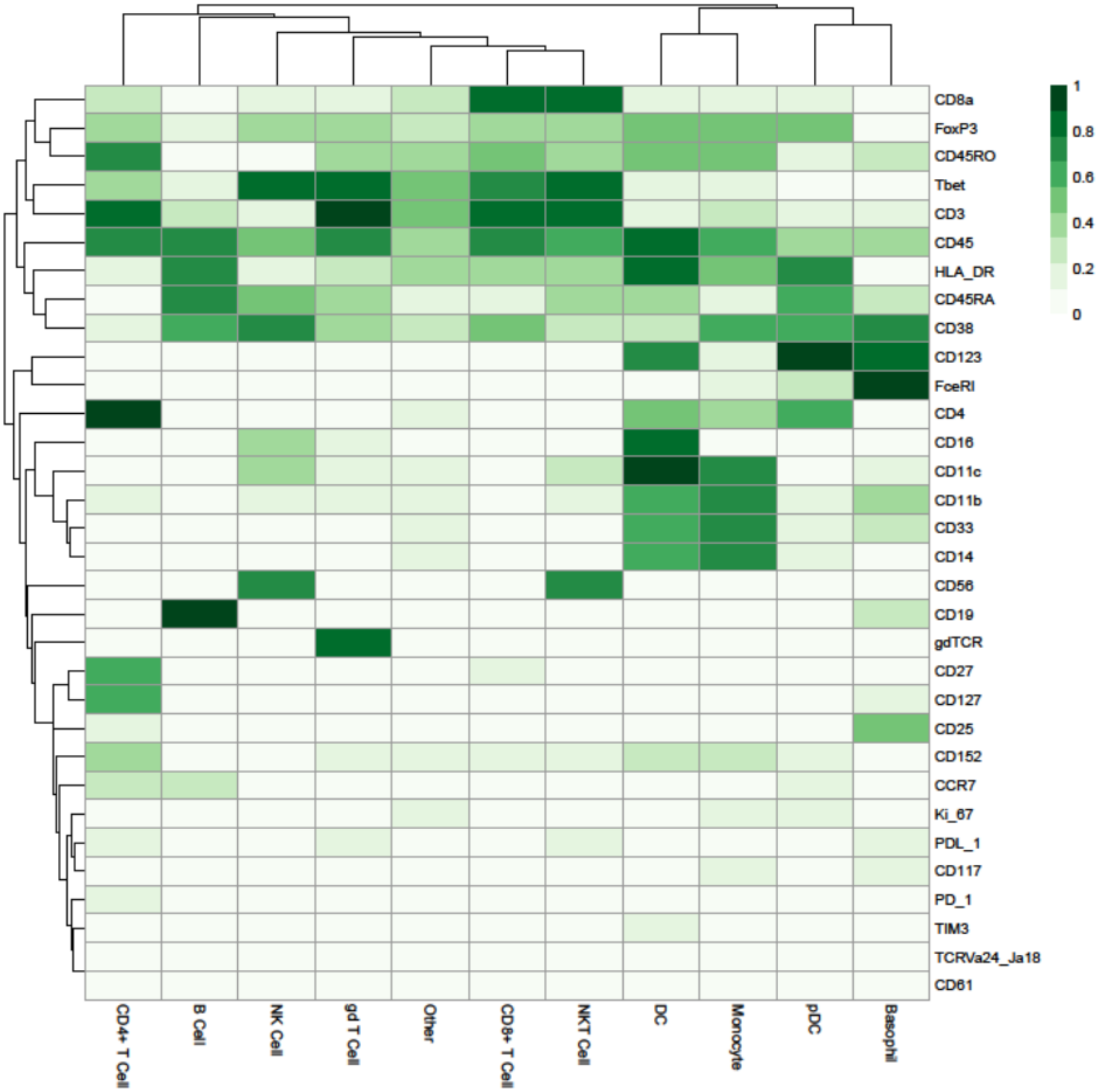

**Figure S9.** Heatmap showing higher-level cluster of cell types.

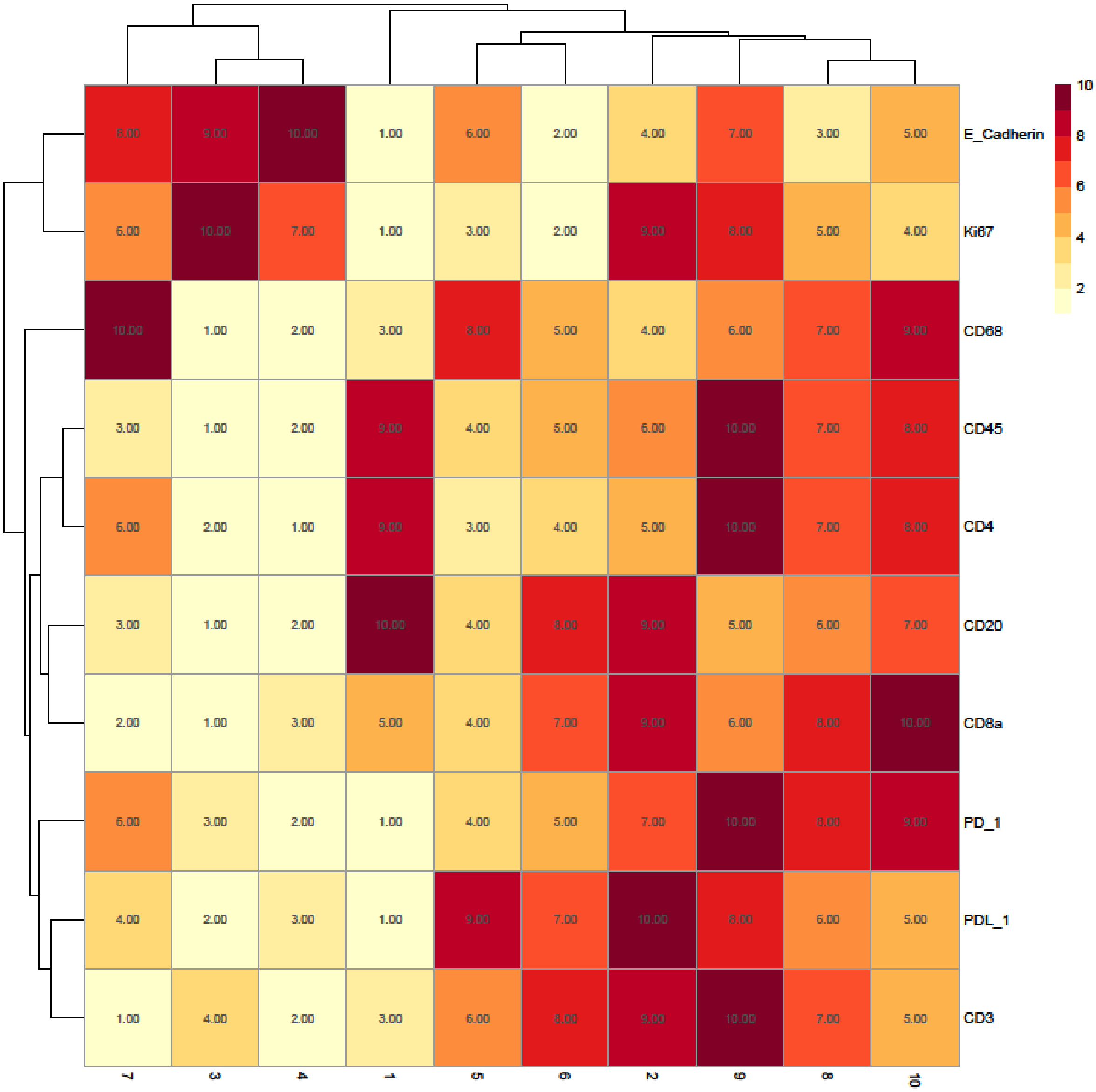

**Figure S10.** Rank heatmap for the head and neck cancer patient IMC data.

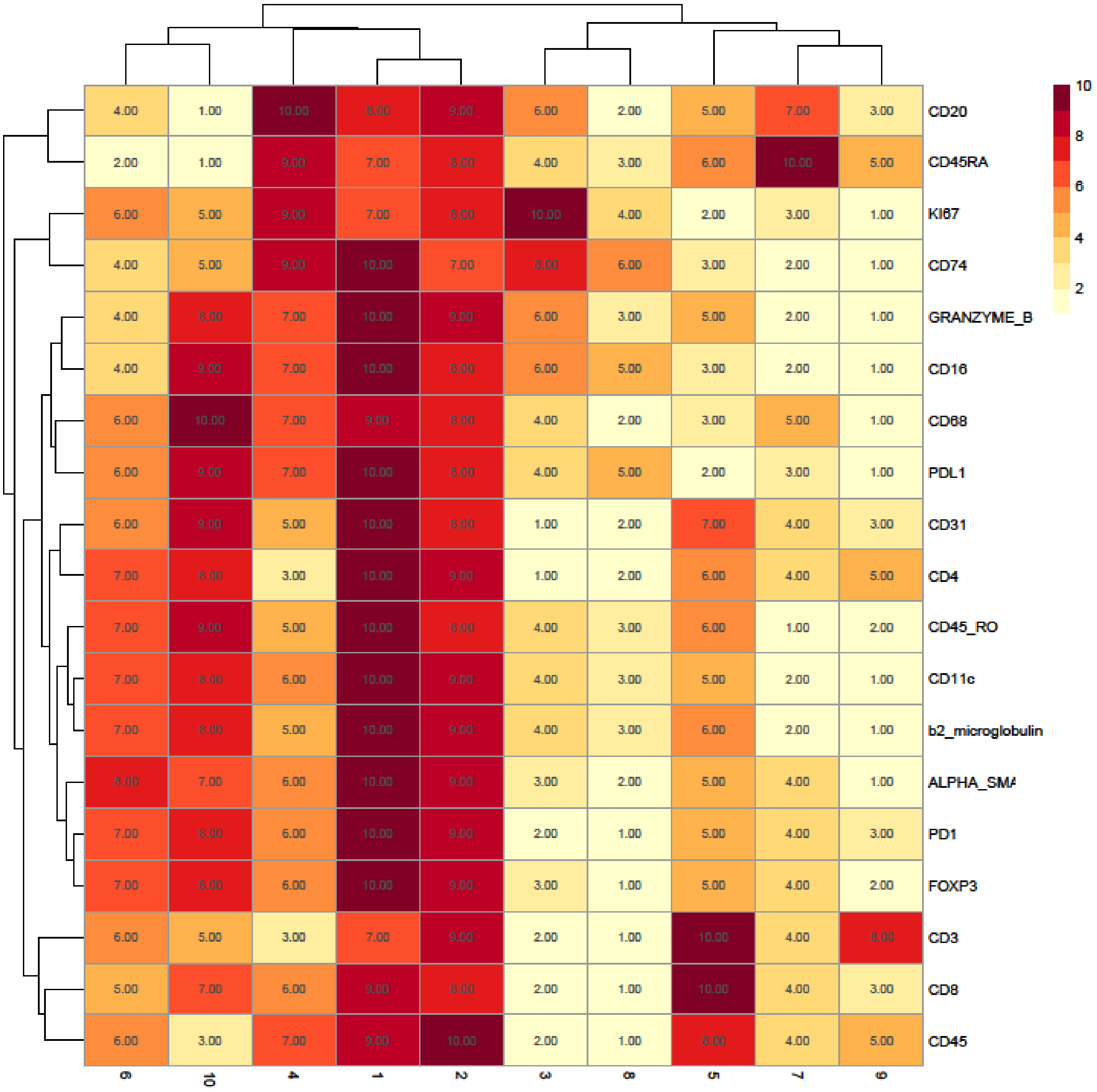

**Figure S11.** Rank heatmap for the diffuse large B cell lymphoma patient IMC data. Clusters 1 and 2 were not used for down-stream analysis as they were deemed to represent minor populations of cells that were non-specifically binding the antibodies as they highly ranked hghly for all markers (including lineage markers) of interest.

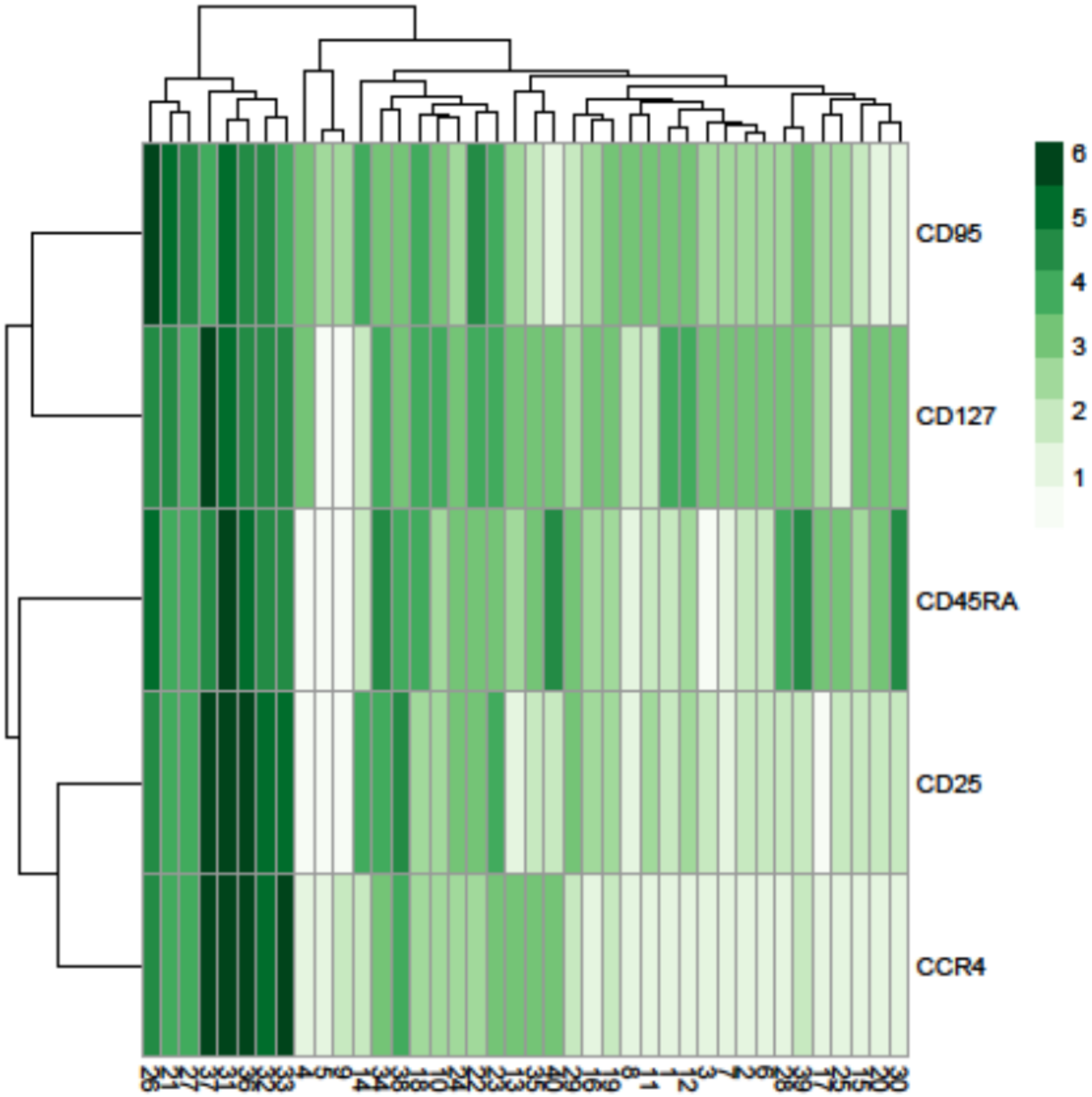

**Figure S12.** Heatmap showing the marker expression of 40 *FlowSOM* clusters. This heatmap was used to identify the cell type of each cluster.

**File S1**

| Equipment | Consumables | Reagents | Reagent Initial Concentration | Reagent Final Concentration |
|---|---|---|---|---|
| Humidified slide chamber (Sigma Aldrich, #H6644-1EA)* | Sterile 3ml pastettes (Alpha Labs, #LW112) | **DPBS,** Ca and Mg free (Gibco, #14190-094) | | |
| Dedicated glassware, coplin jars (Fisher, #10264732) ** | Superfrost Plus™ slides (Fisher, #ES-325) | **Tween-20** (Sigma Aldrich, #P7949-100ml)* | 100% | 0.1% |
| Orbital shaker | Kimberly-Clark™ Precision Wipers (Fisher, #10623111)* | **Antigen retrieval** solution pH 9.0 (R&D Systems, #CTS013) | | 1:10 in milliQ water |
| Water bath | 0.2 syringe filter (TripleRed, #FPE204030) | **Superblock™** (Thermo Fisher, #37515)** | | |
| Fridge | 50ml syringes (Fisher, #10636531) | **Iridium** (Fluidigm, #201192B)*** | 500 µM | 0.5 µM in DPBS |
| Milli-Q water purification system (18.2MΩ.cm) | Filter pipette tips | Metal tagged **antibodies** (Fluidigm) | | Require optimisation for your tissue type |
| Hard slide box for storage in minus 80 (VWR #631-1504) | PAP pen (Abcam, #ab2601) | **FcR block** (Trustain, Biolegend, # 422302) | 100% | 5% |
| Pipettes | | **Xylene** (Acros, #180862500) | | |
| Plastic tweezers, to handle slides *(Farlamedical, #400200)* | | **Ethanol** (Sigma Aldrich, #32221-2.5L) | | |
| Plastic coplin (Thermo Fisher,#E9611) **** | | **BSA** (Sigma Aldrich, #A9647) | 20% To prepare on the same day freshly | 10% |
| | | **Kiovin Hu IgG** Provide by harmacy's hospital | 10% 100mg/ml | 5 mg/ml |
| ** If you don't have one, use a pipette tip box with moist towel at the bottom and piece of parafilm on the tray (this prevents evaporation and stops* | **Normal laboratory handtowels and blue roll are fine to tip excess buffer off slide surface or to place in the bottom of the humidified slide* | **Permeabilization can also be conducted with Triton X100 or saponin* <br><br> *** Alternative blocking solutions* | | |

| | | |
|---|---|---|
| *condensation gathering at the back of the slide, which can pool around edges and dilute antibodies)*<br>***Rinse in water, do not clean with detergent or in laboratory dishwasher (source of barium contamination)*<br>***** Plasticware is required for 96°C water bath, used when dedicated histological autoclave or decloaking chamber is not available; you can also use a 50ml centrifuge tube, loosely capped (Falcon or Corning)* | *chamber – they should not be used to wipe the back of the slides or draw off excess buffer from front – as can transfer lint to the slides.* | *include: BSA, human albumin, human immunoglobulin (KIOVIG from Pharmacy), serum same species as antibodies (mouse serum, Sigma Aldrich, # M5905-10ML).*<br><br>**** DNA stain use alternative histone 3 antibody #3176023D* |

### STAINING PROTOCOL DAY 1: SECTION CUTTING

- Cut the section from the block at 5 µm, Incubation overnight (o/n) at 37°C oven
- Cut a section at 3 µm for Hematoxylin and Eosin (H&E) staining.

### STAINING PROTOCOL DAY 2:

1. *PREPARATION*

- Prepare xylene and ethanol at 50:50 ratio ~50ml of final volume.
- Prepare the following ethanol dilutions, with milli-Q water: 96%, 90%, 80% and 70% (~50ml of final volume for each).
- Dilute the 10X basic antigen retrieval buffer in milli-Q water and heat to 96°C in a water bath, in 50ml falcon tube (leaving the cap loose).
- *BSA (Sigma Aldrich, #A9647): prepare 20%, fresh every time, 0.4g in a plastic tube, add 1ml of superblock solution and incubate in the water bath for 1h, once it is dissolved, top up until 2ml.
- Tween-20 (Sigma Aldrich, #P7949-100ml), prepare the wash buffer at 0.1% in Dulbecco's phosphate buffered saline (DPBS) (500µl of T20 in the DPBS Bottle, filter it every time you use it with 0.2µm filter). stored at room temperature.

2. *Notes for wash steps*

Always conduct washes in in ~50ml of buffer, in coplin jars on an orbital mixer

Never wash in laboratory dishwasher, or with detergent, as this can represent a source of barium contamination. Also, do not use metal slide holders or forceps.

3. STAINING

- Heat the 1X antigen retrieval solution (1:10 dilute in milli-Q water) to 96°C in a water bath.
- Heat a coplin jar and a bottle of 0.1% Tween-20 in DPBS to 37°C in a water bath.
- Prepare the humidified slide chamber (old tip box, with a humid tissue in the bottom and piece of parafilm on the top tray).
- Prepare blocking and antibody incubation buffer on the day of staining – 10% BSA, 0.1% Tween-20, 1/20 dilution of FcR block (Trustain, Biolegend, # 422302) and 1/20 dilution of KIOVIG (Kiovin Hu IgG) in Superblock solution. Use Superblock™ (Thermo Fisher, #37515) to top up to final volume. Avoid mixing, filtering or centrifugation of the Superblock buffer prior to pipetting, let solution settle at 4°C and pipette from top of the solution to avoid any aggregate material that will have sunk to the bottom.

1. *CLEAR WAX FROM SLIDES (You sequentially remove more and more of the wax, with each wash in fresh xylene)*

- Incubate slide 1h at 60ºC for 1h.
- Agitate slides up and down by hand, 3 washes:
- Xylene alone 100% for 5 min
- Xylene alone 100% for 5 min
- Transfer to 50:50 xylene and ethanol mix lifting slides up and down and agitating occasionally for 10 min

> -Xylene washes in the safety cabinet,
>
> -Use 3 separate coplin jars.
>
> -The xylene should be changed regularly, especially the first coplin jar in the sequence, which will have the most wax dissolved in it (readily become cloudy and less efficient).

2. *REHYDRATE TISSUE (outside hood, use plastic tweezers)*

- Remove slides from xylene:ethanol mix in the hood, transfer to 100% ethanol for 5min
- This coplin jar can now be removed from the cabinet:
- Remove slides and transfer to 90% ethanol – mix on orbital shaker for 5 min.
- Remove slides and transfer to 80% ethanol – mix on orbital shaker for 5 min.
- Remove slides and transfer to 70% ethanol – mix on orbital shaker for 5 min.
- Remove slides and transfer to DPBS - mix orbital shaker for 10 min (you cannot reuse this).
- Slide can stay in DPBS until the antigen retrieval solution is ready.

3. *RETRIEVE ANTIGEN*

Fluidigm have validated all their FFPE antibodies using 30 min antigen retrieval, with basic buffer, in a 96°C water bath.

- Heat 1X antigen retrieval solution to 96°C in the water bath (in plastic coplin jar or closed falcon 50ml centrifuge tube).
- When the slides are added the temperature drops down - do not start the timer, until the probe indicates the temperature in the tube reaches 95°C.
- Incubate slides in 1X antigen retrieval solution to 96°C in the water bath (in closed falcon 50ml centrifuge tube) and leave for 30 min.
- Cool down closed falcon under cold running water.
- Transfer slides to DPBS and wash on orbital shaker for 5 min.
- Dry slides with tissue wipers (Thermo Fisher) and leave tot dry for 10 sec.
- Draw a wax ring around the tissue.
- Wash twice with filtered Tween-20 wash buffer on the shaker for 8 min.

(prepare the wash buffer at 0.1% in DPBS (calcium/magnesium free), adding 500µl of T20 in the DPBS bottle, filter it every time you use it with 0.2µm filter and store at room temperature.

- Leave slides in the wash buffer until the block solution is ready.

### *4. BLOCK*

- While sections are in permeabilization buffer (T20 0.1%), prepare sufficient blocking buffer to cover the tissue, within the wax ring.
- In terms of volume – for a very small diameter wax ring 50µl will suffice, a large piece of tissue will require ~250µl.

| Blocking buffer preparation | | | | | | |
|---|---|---|---|---|---|---|
| **Solution** | **Dilute in** | **Initial Concentration** | **Final Concentration** | **Dilution Factor** | **Per 100 µL** | **Per 300 µL** |
| **Tween-20** (Sigma Aldrich, #P7949-100ml)* | Superblock Solution | Prepared first at 10%<br>Then at 1% | 0.1% | 10 | 10 | 30 |
| **Kiovin Hu IgG** | Superblock Solution | 10%<br>100mg/ml | 5mg/ml | 20 | 5 | 15 |
| **FcR block** (Trustain, Biolegend, # 422302) | Superblock Solution | 100% | 5% (1/20) | 20 | 5 | 15 |
| **BSA** (Sigma Aldrich, #A9647) | Superblock Solution | 20%<br>(10g in 100ml=10%) | 10% | 2 | 50 | 150 |
| **Superblock™** (Thermo Fisher, #37515)** | To top up<br>Until your Final Volume | | | | 30 | 90 |

When you add the buffer to the inside of the wax circle, it should be contained and fully cover the tissue.

- Remove slides from DPBS and remove excess buffer by tipping slide.
- Use a lint-free tissue to dry the back of the slide thoroughly and to gently draw excess buffer away from the front, without touching the tissue or wax ring.
- Transfer to humidified box, cover tissue with blocking buffer.
- Incubate at room temperature for 90 min.

### 5. PREPARE ANTIBODY MIX AND INCUBATE

| **Antibody mix** | | | | | | |
|---|---|---|---|---|---|---|
| **Cat#** | **Supplier** | **Tag** | **Target/ Solution** | **Dil Factor (1:X)** | **Initial Conc.** | **Final Conc.** |
| 3170019D | Fluidigm | 170 | CD3 (Polyclonal, C-Terminal) | 400 | | |
| 3156033D | Fluidigm | 156 | CD4 | 100 | | |
| 3152016D | Fluidigm | 152 | CD45 | 1000 | | |
| 3162035D | Fluidigm | 162 | CD8-alpha | 800 | | |
| 3161029D | Fluidigm | 161 | CD20 | 250 | | |
| MAB1561 | R&D Systems | 150 | PD-L1 | 50 | | |
| 3158029D | Fluidigm | 158 | E-CADHERIN | 1000 | | |
| 3159035D | Fluidigm | 159 | CD68 | 400 | | |
| 3165039D | Fluidigm | 165 | PD-1 | 50 | | |
| 3168022D | Fluidigm | 168 | Ki-67 | 400 | | |
| #A9647 | Sigma Aldrich | | BSA Prepare fresh | 2 | 20% | 10% |
| # 422302 | Biolegend | | FCR | 20 | 100% | 5% |
| | Harmacy's hospital | | IgG Hu Kiovig | 20 | 100mg/ml | 5mg/ml |
| #P7949 | Sigma Aldrich | | Tween-20 | 10 | 1% | 0.1% |
| #37515** | Thermo Fisher | | Superblock ™ | To top up, to the required volume | | |

You will require the same volume of antibody mix used for blocking solution and tip off excess blocking buffer onto tissue paper.

Use a lint-free tissue to dry the back of the slide thoroughly and to gently draw excess buffer away from the front, without touching the tissue or wax ring.

- Incubate slide o/n with antibody mix at 4$^{o}$C in humidified chamber.

### STAINING PROTOCOL DAY 3:

1. *WASH IN PERMEABLIZATION BUFFER*

- Tip off the antibody mix onto tissue paper.
- Flush the slide surface thoroughly and gently with a pastette and permebalization buffer (DPBS with added 0.1% tween) at room temperature.
- Wash in the same room temperature buffer, 50ml in coplin jar, on the mixer for 8 min.
- Repeat wash for another 8 min.

2. *WASH IN DPBS*

- In the same Jar, decant T20 and add DPBS wash buffer at room temperature, using around 50ml in a clean coplin jar and then place on the mixer for 8 min.
- Repeat wash for another 8 min.

3. *DNA STAIN*

For initial testing, prepare iridium (Fluidigm) in DPBS at a final concentration of 0.5 µM (may require some optimization for your tissue type).

Dilute 125µM iridium stock (Fluidigm 201192A) 1/250 with DPBS.

- Tip off excess DPBS onto tissue paper.
- Use a lint-free tissue to dry the back of the slide thoroughly and to gently draw excess buffer away from the front, without touching the tissue or wax ring.
- Place slide in humidified chamber and load iridium solution into the wax ring so that it covers the tissue.
- Incubate at room temperature for 30 min.
- Flush the slide surface thoroughly with room temperature DPBS gently using a pastette.
- Wash in room temperature DPBS on the mixer for 5 min.
- Wash in Milli-Q, in 50ml in a clean coplin jar, on the mixer for 5 min.
- Air dry.

Slides can be stored for several months at room temperature in in a cool, dry, dust-free storage container.

### File S2

**Protocol for sectioning and staining of diffuse large B-cell lymphoma lymph node section**

1. A 5μm tissue section was deparaffinized with xylene and sequentially rehydrated in graded ethanol. Heat-induced antigen retrieval was performed in an automated pressure cooker (Menapath Antigen Access Unit, Menarini) at 125$^{o}$C for 2 min in Antigen Retrieval Reagent-Basic (R&D Systems).
2. The tissue was then permeabilized with Dulbecco's PBS (DPBS), 0.1% Tween-20 for 15 mins and blocked with Superblock (Thermo Fisher Scientific), 0.1% Tween-20, Fc Receptor Blocking Solution (1:20) (Biolegend) (blocking buffer) for 2 h at RT.
3. Antibodies were diluted together as according to Table S3 in blocking buffer and applied to the tissue overnight at 4 $^{o}$C.
4. After washing, the slide was incubated with the DNA intercalator (Cell-ID Intercalator-Ir, Fluidigm) for 30 min at RT.
5. The slide was then briefly washed with $H_2O$ and air dried.
6. Acquisition of 1 mm$^2$ tissue region was carried out using the Hyperion imaging system (Fluidigm) at 200 Hz, with resolution of 1 μm/pixel.
